# Tau interactome mapping reveals dynamic processes in synapses and mitochondria associated with neurodegenerative disease

**DOI:** 10.1101/2021.06.17.448349

**Authors:** Tara E. Tracy, Jesus Madero-Pérez, Danielle Swaney, Timothy S. Chang, Michelle Moritz, Csaba Konrad, Michael E. Ward, Erica Stevenson, Ruth Hüttenhain, Grant Kauwe, Maria Mercedes, Lauren Sweetland-Martin, Xu Chen, Sue-Ann Mok, Maria Telpoukhovskaia, Sang-Won Min, Chao Wang, Peter Dongmin Sohn, Jordie Martin, Yungui Zhou, Giovanni Manfredi, Giovanni Coppola, Nevan J. Krogan, Daniel H. Geschwind, Li Gan

**Author notes:** Equal contributions (Tara E. Tracy, Jesus Madero-Pérez, Danielle Swaney).

## Abstract

Tau (MAPT) drives neuronal dysfunction in Alzheimer’s disease (AD) and other tauopathies. To dissect the underlying mechanisms, we combined an engineered ascorbic acid peroxidase (APEX) approach with quantitative affinity purification mass spectrometry (AP-MS) followed by proximity ligation assay (PLA) to characterize Tau interactomes modified by neuronal activity and mutations that cause frontotemporal dementia (FTD) in human induced pluripotent stem cell (iPSC)-derived neurons. We established activity-dependent interactions of Tau with presynaptic vesicle proteins during Tau secretion and mapped the exact APEX-tau-induced biotinylated tyrosines to the cytosolic domains of the interacting vesicular proteins. We showed that FTD mutations impair bioenergetics and markedly diminished Tau’s interaction with mitochondria proteins, which were downregulated in AD brains of multiple cohorts and correlated with disease severity. These multi-modal and dynamic Tau interactomes with unprecedented spatiotemporal resolution shed novel insights into Tau’s role in neuronal function and disease-related processes with potential therapeutic targets to block Tau-mediated pathogenesis.

## INTRODUCTION

Accumulation of pathological Tau protein in the brain is associated with cognitive decline in Alzheimer’s disease (AD) and other neurodegenerative diseases classified as tauopathies. Mutations in the microtubule-associated protein Tau (*MAPT)* gene are sufficient to cause familial frontotemporal dementia (FTD) and Tau is the major factor that drives neurodegeneration in other primary tauopathies including corticobasal syndrome and progressive supranuclear palsy syndrome. In healthy neurons, Tau has an intrinsically disordered structure and interacts preferentially with microtubules to provide stabilization (Elie et al., 2015; Santarella et al., 2004). Tau undergoes extensive posttranslational modifications including phosphorylation and acetylation, which are modified by disease (Min et al., 2015; Morris et al., 2015; Wesseling et al., 2020). During this pathogenic process, Tau becomes toxic to neurons and impinges on various processes important for neuronal physiology including cytoskeletal dynamics (Bardai et al., 2018; DuBoff et al., 2012), axonal transport (Ishihara et al., 1999; Ittner et al., 2008), synaptic transmission (Hoover et al., 2010; Ittner et al., 2010; McInnes et al., 2018; Tracy et al., 2016; Yoshiyama et al., 2007), proteostasis (Myeku et al., 2016), nuclear transport (Eftekharzadeh et al., 2018), and axon initial segment function (Li et al., 2011; Sohn et al., 2019; Sohn et al., 2016). Direct interactions of Tau with specific proteins or protein complexes have been identified that can alter the aforementioned cellular processes (Eftekharzadeh et al., 2018; Ittner et al., 2010; Ittner et al., 2009; McInnes et al., 2018; Sohn et al., 2019; Vanderweyde et al., 2016), indicating that Tau can have a significant impact on neuron function through its microtubule-independent protein-protein interactions (Morris et al., 2011). Affinity purification mass spectrometry (AP-MS)-based characterizations of the Tau interactome have focused on mouse neurons in wild-type mice (Liu et al., 2016; Wang et al., 2017b), or tauopathy mice that express high levels of mutant Tau (Choi et al., 2020; Maziuk et al., 2018). Other studies examined Tau interactomes in SH-SY5Y neuroblastoma cells (Gunawardana et al., 2015), or a mixed population of neuroprogenitor-derived human ReN cells (Wang et al., 2019). However, none of the studies have directly compared the interactomes of wild-type Tau and FTD mutant Tau or captured how the Tau interactome is affected by neuronal activity in human neurons.

Neuronal activity triggers the secretion of Tau from neurons (Pooler et al., 2013; Wu et al., 2016; Yamada et al., 2014) and the translocation of Tau into synapses (Frandemiche et al., 2014). Secreted Tau can spread trans-neuronally between synaptically connected neurons (Wang et al., 2017c; Wu et al., 2016), and indeed pathological analyses from postmortem human tauopathy brains suggest that Tau spreads across functionally connected regions of the brain (Vogel et al., 2020). Depolarization of human AD brain synaptosome fractions triggers Tau release from presynaptic terminals (Sokolow et al., 2015), suggesting that Tau secretion is regulated by presynaptic function and that yet unknown Tau protein-protein interactions at synapses may be involved.

To characterize the Tau interactome in human neurons, we took advantage of both engineered ascorbic acid peroxidase (APEX) proximity-dependent mapping with mass spectrometry and AP-MS to reveal the dynamic and multifaceted interactome of Tau in human induced pluripotent stem cell (iPSC)-derived glutamatergic neurons. The APEX technique has been used to generate precise spatial and temporal maps of subcellular compartment proteomes (Loh et al., 2016; Rhee et al., 2013) and protein localization during intracellular trafficking (Lobingier et al., 2017). Similarly, APEX mapping enabled us to elucidate the precise spatial and temporal regulation of the Tau interactome during enhanced neuronal activity. Furthermore AP-MS allowed us to quantitatively compare protein-protein interactions of wild-type Tau with that of FTD mutant Tau in human neurons. We identified known Tau interactors as well as novel interacting proteins that could mediate disease pathogenesis. Importantly, we show that the levels of key Tau-interacting proteins were altered in human AD brain samples. Together, the APEX and AP-MS analyses of the Tau interactome provide a comprehensive mapping of protein-protein interactions which could be targeted therapeutically to slow Tau-mediated disease progression.

## RESULTS

### Labeling of the Tau interactome in living human iPSC-derived neurons

Expression of the transcription factor Neurogenin-2 (NGN2) in human iPSCs drives them into a highly homogenous culture of functional glutamatergic neurons (Zhang et al., 2013). We previously developed a platform called i^3^Neurons via genetic editing of a human iPSC line (WTC11) to carry inducible expression of NGN2, allowing a greatly simplified, scalable and homogeneous differentiation into neurons (Wang et al., 2017a). To investigate Tau interactions in living neurons, we used a similar approach to engineer i^3^Neurons that had one copy of the doxycycline-inducible NGN2 transgene integrated into one allele of the AAVS1 safe harbor locus, while the other allele was integrated with a transgene for doxycycline-inducible expression of human Tau tagged with a flag epitope and APEX2, a highly active version of the engineered ascorbate peroxidase (Lam et al., 2015) (Fig. 1A). Induction of Cre/loxP recombination excised the puromycin resistance gene in iPSCs within the NGN2 transgene cassette to allow for puromycin selection following another round of genome editing. Targeted editing of the second AAVS1 allele by TALENs was followed by puromycin selection to isolate iPSC clones carrying the human Tau transgene. Proteolytic cleavage of Tau has been reported in cultured human neuron models and the accumulation of Tau fragments is associated with neurodegenerative phenotypes (Ehrlich et al., 2015; Fong et al., 2013). To capture the proteins interacting with N- or C-terminus truncated Tau, we generated two independent isogenic clonal iPSC lines expressing either the N-terminus- or the C-terminus-tagged wild-type 2N4R Tau (Tau) tagged with APEX2 and flag epitope (Figure 1B). As a control for Tau-specific interactions we also made an isogenic iPSC clone with expression of APEX2-tagged Tubulin α 1b (APEX-α Tubulin). All iPSC clones exhibit normal karyotypes (Figure S1).

**Figure 1.**
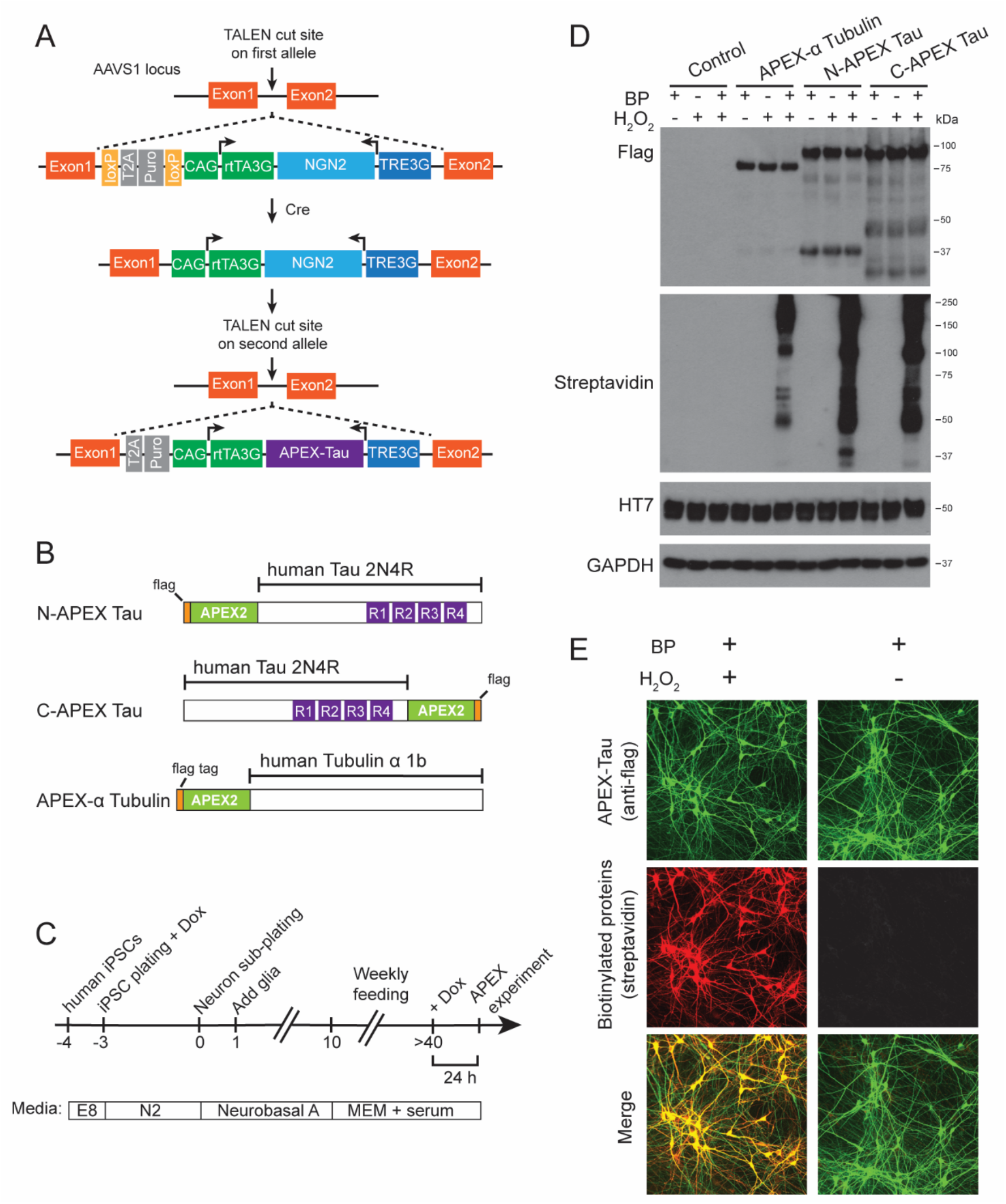
APEX-Tau mediated proximity-dependent biotinylation in human iPSC-derived neurons. (A) Diagram illustrating the TALEN-mediated targeted integration of the NGN2 transgene into one allele of the AAVS1 locus in human iPSCs, followed by the integration of the APEX-Tau transgene into the second AAVS1 allele. (B) Design of the APEX-flag tagged full-length human Tau (2N4R) and Tubulin α 1b constructs each integrated into a stable iPSC line. See also Figure S1. (C) Timeline and workflow of i^3^Neuron differentiation and experimental analyses. (D) Western blot analysis of N- and C-terminal APEX tagged Tau and APEX-Tubulin expressed in i^3^Neurons. Biotinylated proteins were only detected in lysates from neurons that were treated with 30 minutes of biotin-phenol (BP) and 1 min of H_2_O_2_ to stimulate APEX activity. (E) Representative confocal images of APEX-Tau expression (green) and biotinylated proteins (red) in i^3^Neurons. Biotinylation of proteins in APEX-Tau neurons was triggered by the 1 min treatment with H_2_O_2_.

After the induction of neuronal differentiation and maturation, neurons were treated with doxycycline at 6 weeks to turn on the expression of APEX2-tagged Tau or APEX-α Tubulin for 24 hours before experiments (Figure 1C). Immunoblotting revealed same levels of full-length N-APEX Tau or C-APEX Tau proteins with some distinct N- and C-terminus fragments labeled by the flag epitope (Figure 1D). Biotin-phenol (BP) treatment and a brief H_2_O_2_ exposure induced biotinylation of proteins associated with α Tubulin and with N-APEX- and C-APEX Tau at similar levels (Figure 1D). The H_2_O_2_-dependent biotinylation of proteins was strongly co-localized with APEX2-tagged Tau in the soma and in neuronal processes (Figure 1E).

### Spatially defined proteomic mapping of the Tau interactome

Human iPSC-derived neurons expressing N- or C-terminal tagged APEX-Tau were treated with BP, followed by 1 min of H_2_O_2_ to stimulate biotinylation of Tau-associated proteins in a snapshot of time. An anti-biotin antibody was then used to enrich for biotin-modified peptides (Udeshi et al., 2017), and this enriched population was detected by mass spectrometry enabling direct detection of biotinylated tyrosine residues in close proximity to APEX-Tau (Figure 2A). Proteins biotinylated by N-APEX Tau, C-APEX Tau or APEX-α Tubulin were defined as those enriched in the majority of neuronal cultures expressing those constructs (Table S1).

**Figure 2.**
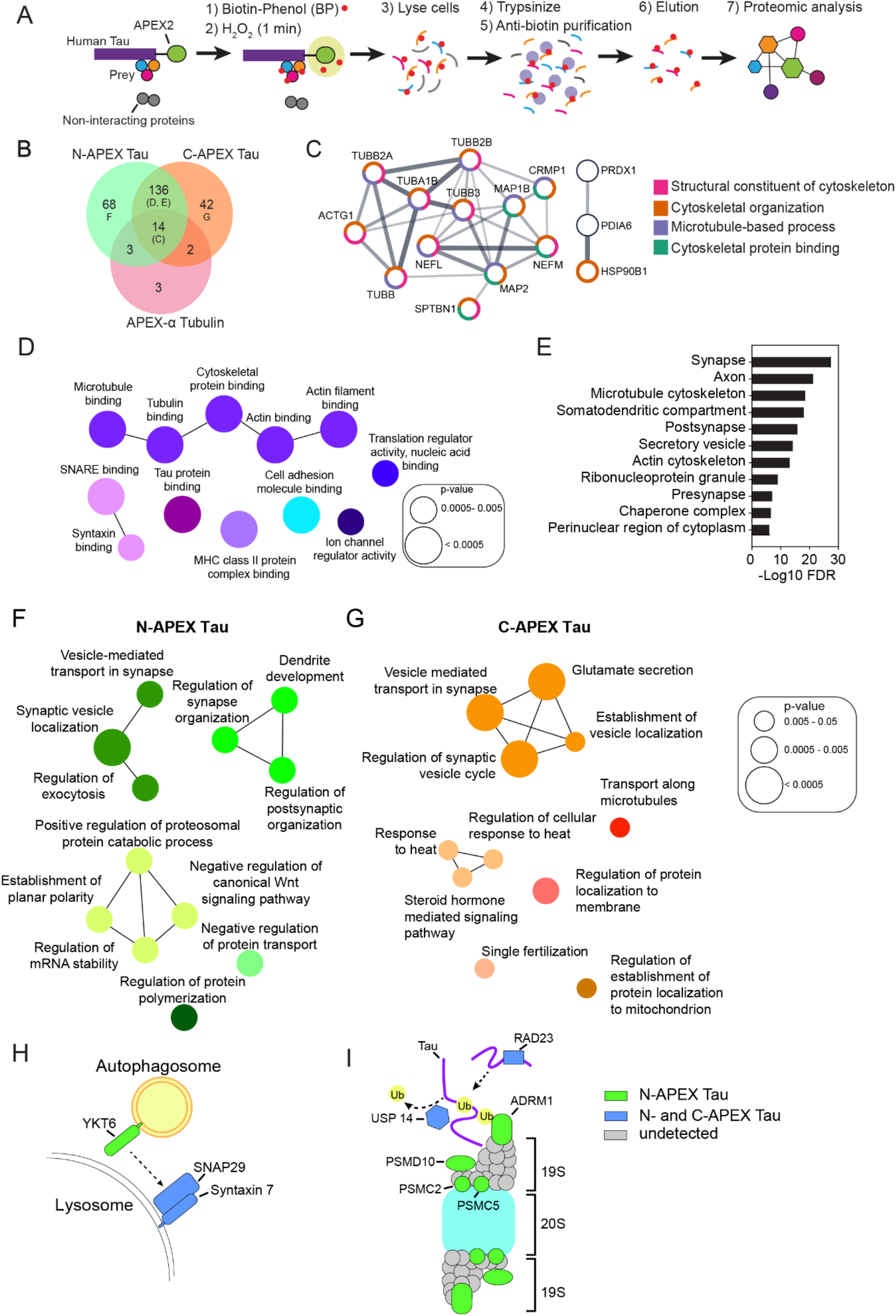
Subcellular and subprotein domain Tau interactome in living human neurons identified by APEX-mediated biotinylation. (**A**) Schematic of the workflow for the proximity-dependent identification of Tau-associated proteins by mass spectrometry. (**B**) Venn diagram of the biotinylated proteins detected in N-APEX-Tau, C-APEX-Tau and APEX-Tubulin expressing human neurons. See also Table S1. (**C**) Proteins biotinylated by N-APEX-Tau, C-APEX-Tau and APEX-Tubulin include components of the microtubule cytoskeleton network. Only interconnected nodes based on the STRING database are shown. The category of cytoskeletal function by pathway analyses is grouped by color. (**D**) ClueGO molecular function pathway enrichment of proteins biotinylated by both N-APEX-Tau and C-APEX-Tau but not APEX-Tubulin. Node colors denote functionally grouped networks (kappa connectivity score ≥ 30%). (**E**) Graph of Gene Set Enrichment Analysis (GSEA) on proteins biotinylated by both N-APEX- and C-APEX-Tau showing the major cellular component categories identified. (**F, G**) ClueGO biological processes pathway enrichment of proteins biotinylated only by N-APEX-Tau (**F**) and only by C-APEX-Tau (**G**). (**H, I**) Illustrations of sub-protein level APEX-Tau associations identified that regulate autophagy (**H**) and proteasome degradation (**I**).

To identify Tau-specific interacting proteins, we compared the biotinylated proteins between the N-APEX Tau and C-APEX Tau neurons and the APEX-α Tubulin neurons (Figure 2B). The majority of proteins enriched for association with α Tubulin were also detected in the Tau interactome results, consistent with the close interaction of Tau and Tubulin on microtubules. Indeed, pathway analyses revealed that most of the proteins associated with both α Tubulin and Tau were components of microtubules and other cytoskeletal-binding proteins (Figure 2C), demonstrating the specific enrichment for proteins in close proximity with high spatial resolution using this method. To focus our analyses on Tau-specific interactions, we removed the 19 proteins co-associated with both Tau and α Tubulin from further analyses. Of the 246 putative Tau-interacting proteins, 136 were enriched in both N-APEX and C-APEX Tau neurons (Figure 2B). Gene Ontology (GO) enrichment analyses revealed that the molecular functions of these overlapping proteins largely comprised microtubule and actin cytoskeletal networks (Figure 2D), as well as previously established Tau-binding proteins including α-Synuclein (SNCA) (Benussi et al., 2005), MAP1A (Alonso et al., 1997), HSP70 (Taylor et al., 2018), HSP90 (Weickert et al., 2020) and Protein Phosphatase 5 (PPC5C), which forms a complex with HSP90 (Silverstein et al., 1997) and dephosphorylates Tau (Liu et al., 2005) (Table S1). In addition, the snapshot of the interactome revealed an enrichment for components of SNARE complexes including SNAP29, Syntaxin 7 (STX7), SNAP25, ɣ-SNAP, and Munc18 (STXBP1), evidence of Tau’s localization at sites of membrane and vesicle fusion. Major classifications of biotinylated proteins by cellular components included synapses, microtubule an actin cytoskeletons, secretory vesicles, and ribonucleoprotein granules (Figure 2E).

We next examined the differential N- and C-terminal interactomes of Tau. Strikingly, 45% of all interactions were uniquely detected in either N-APEX or C-APEX Tau neurons (Figure 2B). Enrichment analyses of the distinct N- and C-terminal Tau associated proteins revealed networks involved in synaptic vesicle regulation (Figures 2F and 2G). While N-APEX Tau biotinylated proteins are mostly involved in active zone docking including Dynamin 1 (DNM1), α-/β-SNAP (NAPA/B), RAB3GAP1, and Liprin α3 (PPFIA3), the C-APEX Tau interacting proteins are important for the fusion of vesicles, such as Syntaxin 1A/1B (STX1A/1B), RAB3A, RIMS1 and Mint1 (APBA1). These findings showed that Tau is localized at sites of presynaptic vesicle fusion and could have domain-specific interactions with synaptic vesicle associated proteins (Table S1). Interestingly, the N-APEX Tau interactome also included proteins that regulate dendritic development and postsynaptic organization including ABI2, Cadherin-2 (CDH2), Nectin (PVRL1), PAK3, 14-3-3ζ (YWHAZ), and GRIP1, indicating that the N-terminus of Tau could impact both pre- and post-synaptic structures through multiple pathways (Figure 2F).

Tau interactomes also included proteins that modulate protein degradation. The N-APEX Tau interactome included YKT6, a SNARE protein on autophagosomes critical for lysosomal fusion (Matsui et al., 2018). For membrane fusion with the lysosome to occur, YKT6 forms a SNARE complex with SNAP29 and lysosomal Syntaxin 7, which were biotinylated by both N- and C-APEX Tau (Figure 2H), suggesting that Tau is localized to sites of autophagosome-lysosome fusion. Moreover, we found that a regulatory component of the 19S cap complex of the proteosome, ADRM1, and three 19S subunits of the 26S proteasome—PSMC2, PSMC5 and PSMD10—were only enriched in the N-terminal Tau interactome (Figure 2I). This spatially-restricted capture of the 19S cap components suggests that the N-terminus is key for the interaction of Tau with the 26S proteasome in human neurons. In contrast, USP14 and RAD23, involved in the ubiquitin-dependent proteolysis of Tau by proteasome, were identified in both N- and C-APEX Tau interactomes, suggesting that the interaction of Tau with ancillary factors involved in proteasome function is not domain-specific.

### Biotinylation sites reveal Tau-interacting proteins at subcellular and amino acid levels

**T**he anti-biotin purification strategy enabled the direct detection of >600 distinct biotinylated tyrosine residues on Tau- and tubulin-associated proteins (Figures 3A and 3B, Table S2). We observed that the APEX-α Tubulin, N-APEX-Tau, and C-APEX-Tau biotinylated largely the exact sites on α Tubulin, β Tubulin, MAP1B, and MAP2, supporting the close structural proximity of these microtubule cytoskeletal proteins (Figure 3C). For full length Tau (2N4R), four out of the five tyrosine residues were biotinylated by both N- and C-terminal tagged APEX; one in the N-terminus was only biotinylated in N-APEX Tau neurons (Figure 3C).

**Figure 3.**
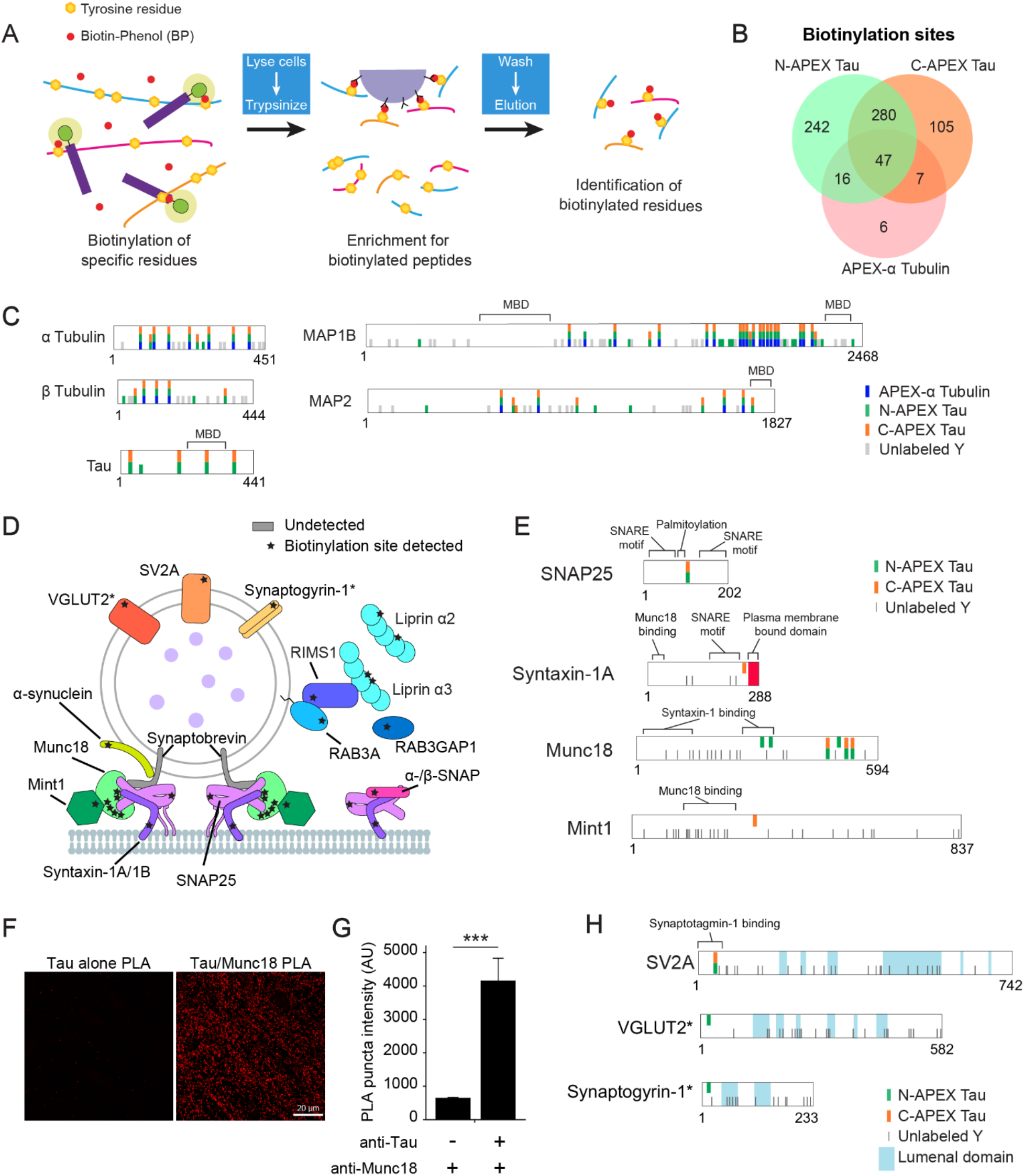
Mapping of the biotinylation sites on Tau-associated proteins. (**A**) Diagram of the workflow used to enrich and identify biotinylated peptides following the anti-biotin pulldown of proteins biotinylated by APEX-Tau. (**B**) Venn analysis of the biotinylated tyrosines on peptides detected in human neurons expressing N-APEX-Tau, C-APEX-Tau or APEX-Tubulin. See also Table S2. (**C**) Representations of the biotinylated and unlabeled tyrosines (Y) on cytoskeletal proteins biotinylated by APEX-Tubulin, N- and C-APEX Tau. See also Figure S2. (**D**) Illustration showing the biotinylation sites (stars) on SNARE complex and synaptic vesicle-associated proteins that were labeled by APEX-Tau. Biotinylation was detected on the t-SNAREs, Syntaxin and SNAP25, however the v-SNARE, Synaptobrevin, was not detected. Biotinylation was also detected on the cytosolic domains of integral vesicle membrane proteins rather than on their lumenal domains. (**E**) Tyrosine residues biotinylated by N-and C-APEX Tau mapped onto SNARE complex proteins adjacent to structural and functional domains that mediate vesicle fusion. (**F**) Representative images of the PLA reaction (red) in human neurons with the Tau5 antibody alone (left) and with both the Tau5 and Munc18 antibodies (right). (**G**) Quantification of PLA puncta fluorescence intensity in human neurons with and without the Munc18 antibody (n= 6 images/group, ***p < 0.001, unpaired student’s *t*-test). Values are means ± SEM. See also Figure S3. (**H**) Locations of biotinylated residues detected in relation to the topology of integral synaptic vesicle membrane proteins. Biotinylation sites were detected on the cytosolic, not the lumenal domains. Asterisks denote biotinylated proteins that were detected in less than half of the samples (VGLUT2 n= 3 and Synaptogyrin-1 n= 2 out of 7 samples).

Nuclear localization of Tau was supported by the detection of four biotinylated residues on Emerin (EMD) (Figure S2A), an integral component of the inner nuclear membrane which functions in the nuclear lamina (Berk et al., 2013) as well as RUVBL1 and RUVBL2 (Table S2), homologous nuclear proteins critical for chromatin-remodeling and transcription (Jha et al., 2008; Jonsson et al., 2004). Specific biotinylation sites were also detected on RNA- and DNA-binding proteins (Figure S2B). Consistent with the perinuclear cytosolic association of Tau and Importin-β (KPNB1) (Lee et al., 2005; Zachariae and Grubmuller, 2008), both N- and C-APEX Tau biotinylated Importin-β within the RanGTP binding site (Figure S2A), and N-APEX Tau biotinylated RANGAP1 which promotes the release of RanGTP from Importin-β when it returns to the cytosol from the nucleus.

Domain-specific biotinylation sites on SNARE complex and synaptic vesicle proteins were detected (Figure 3D). We captured the known interaction between Tau and α-Synuclein (Giasson et al., 2003), by detecting biotinylated tyrosine 39 in the N-terminal domain of α-Synuclein (Table S2). Biotinylation sites were detected on two of the three components in the assembled SNARE complex required for synaptic vesicle fusion, Syntaxin-1A/1B and SNAP25 (Figure 3E). Six biotinylation sites were detected on Munc18 (Figure 3E), a protein essential for vesicle fusion that interacts directly with the SNARE complex via Syntaxin-1 (Dulubova et al., 2007). The recruitment of Munc18 to sites of vesicle release is facilitated by its interaction with Mint1 (Biederer and Sudhof, 2000), which was also biotinylated by C-APEX-Tau. In the human iPSC-derived neurons, we confirmed that Munc18 co-localizes with Synapsin at presynaptic terminals (Figure S3A). Using a proximity ligation assay (PLA) with primary antibodies against Tau (Tau5) and Munc18 to examine the association with endogenous human Tau, we found that the PLA signal was significantly higher in neurons stained with the Tau5 and Munc18 antibodies compared to a Tau5 antibody alone control (Figure 3F, G), supporting the close proximity (<40nm) of endogenous Tau and Munc18. The only biotinylation site on SV2A, a synaptic vesicle integral membrane protein, was on a cytosolic domain of the protein (Figure 3H). Two additional vesicular proteins, Synaptogyrin 1 (SYNGR1) and VGLUT2 (SLC17A6), were biotinylated on their cytosolic domains by N-APEX Tau (Figure 3H). These results suggest that Tau is closely associated with the cytosolic membrane surface, as opposed to the lumenal membrane surface, of synaptic vesicles with assembled SNARE complexes at vesicle fusion sites in human neurons.

### Mapping the activity-dependent change in the Tau interactome

We next used APEX-Tau to obtain a snapshot of the dynamic changes in the Tau interactome that occur when Tau is released from neurons during enhanced neuronal activity. Consistent with previous reports in mammalian neurons (Pooler et al., 2013; Schoch et al., 2016; Yamada et al., 2014), increasing the excitability of human iPSC-derived neurons with a 30-minute treatment of 50mM KCl caused a significant rise in extracellular Tau levels that was dependent on intracellular calcium (Figure 4A), but did not promote cell death (Figure 4B). To detect the changes in the Tau interactome during enhanced neuronal activity, we performed APEX reactions and anti-biotin enrichment for both N- and C-APEX Tau as described earlier (Figure 2A), except that neurons were treated with 50mM KCl for 30 minutes prior to cell harvest. The biotinylated proteins enriched in neuronal cultures treated with 50mM KCl (Table S3) were compared to the biotinylated proteins enriched in unstimulated neuronal cultures (Figure 2B, Table S1). Most of the biotinylated proteins identified, 76% in N-APEX Tau neurons and 68% in C-APEX Tau neurons, were detected in both unstimulated and 50mM KCl conditions (Figure 4C, Table S3), suggesting that the majority of the Tau interactome, including many cytoskeletal-associated proteins, did not change in response to activity. Of the remaining biotinylated proteins affected by activity (Figure 4C, Table S3), most were detected only in neurons treated with 50mM KCl (17% and 28% in N-APEX and C-APEX Tau neurons, respectively) with fewer biotinylated proteins detected exclusively in unstimulated neurons (6% and 4% in N-APEX and C-APEX Tau neurons, respectively).

**Figure 4.**
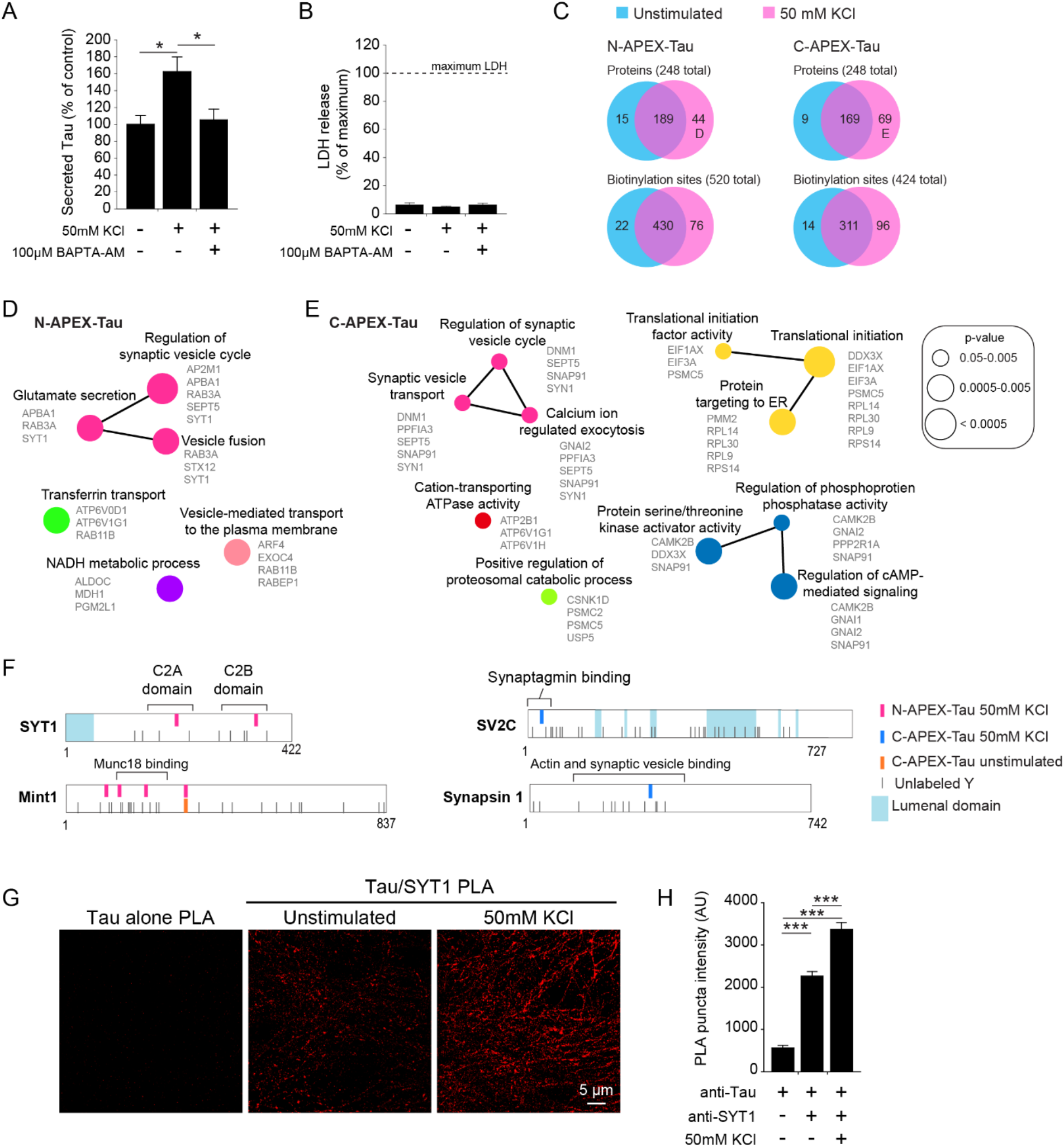
Activity-induced changes in the Tau interactome during activity-dependent Tau secretion from human neurons. (**A**) Quantification of Tau secreted from human neurons during 30 minutes with or without treatment with high KCl to enhance neuronal activity and BAPTA-AM, a membrane permeable calcium chelator. Levels of secreted Tau measured by ELISA were normalized to total Tau levels in the neuron culture (n= 8-9 cultures/group, *p < 0.05, one-way ANOVA, Bonferroni *post*-hoc analyses). Values are means ± SEM. (**B**) Lactate dehydrogenase (LDH) levels were measured in the media collected from human neurons after 30 minutes with or without high KCl treatment (n= 8-9 cultures/group). (**C**) Venn analyses of the biotinylated proteins and the individual biotinylation sites that were labeled by N-APEX Tau and C-APEX Tau without stimulation (blue, data from Figure 2B) and with enhanced neuronal activity (red). Human neurons were either unstimulated or treated with 50 mM KCl for 30 minutes followed by stimulation of APEX activity to induce the biotinylation of Tau interacting proteins. See also Table S3. (**D, E**) ClueGO biological processes pathway enrichment of proteins biotinylated by (**D**) N-APEX Tau or by (**E**) C-APEX-Tau in neurons with KCl-induced increased activity. These analyses were performed on the 44 (**D**) and 70 (**E**) biotinylated proteins labeled in panel **C**. (**F**) Activity-induced biotinylated residues were detected on synaptic vesicle-associated proteins. (**G**) Representative confocal images of PLA fluorescence with Tau5 and Synaptotagmin-1 antibodies in human neurons that were unstimulated or stimulated with 50mM KCl for 30 minutes. (**H**) PLA puncta fluorescence intensity was quantified in human neurons stained with the Tau5 and Synaptotagmin-1 antibodies with and without enhanced activity in high KCl (n= 6-12 images/group, *** p < 0.001, one-way ANOVA, Bonferroni *post*-hoc analyses). Values are means ± SEM. See also Figure S3.

To characterize the proteins that interact with Tau during activity-induced Tau secretion, we performed network analyses on the biotinylated proteins that were enriched only in the neurons treated with 50mM KCl. This revealed activity-dependent networks that regulate synaptic vesicle exocytosis which were common to both N-APEX and C-APEX Tau-expressing neurons (Figures 4D and 4E). Activity-dependent biotinylated residues were detected on SNARE complex-associated proteins Synaptotagmin 1 (SYT1) and Mint1, and synaptic vesicle proteins, SV2C and Synapsin 1 (SYN1) (Figure 4F, Table S3). Further supporting the interaction of Tau with the cytosolic side of synaptic vesicle membranes, the biotinylated residues in Synaptotagmin 1 and SV2C were in the cytosolic, rather than lumenal, domains.

Two activity-dependent N-APEX Tau-mediated biotinylation sites were detected within the calcium-binding domains of Synaptotagmin-1, which is a vesicular protein that acts as a calcium sensor and triggers synaptic vesicle fusion at the presynaptic terminal. The localization of Synaptotagmin-1 to presynaptic terminals of human iPSC-derived neurons was confirmed by co-immunostaining for the presynaptic marker Synapsin 1 (Figure S3B). The intensity of the PLA signal with Tau and Synaptotagmin-1 antibodies was significantly increased in neurons following treatment with 50mM KCl compared to unstimulated neurons (Figure 4G and 4H). These data support a model in which Tau associates with proteins in SNARE complexes and synaptic vesicles at the presynaptic active zone, and indicate that activity-dependent Tau secretion from neurons could involve enhanced interaction with Synaptotagmin-1 during the fusion of synaptic vesicle at the presynaptic membrane.

### Familial FTD mutations modify the Tau interactome

Tau mutations cause FTD. To identify the steady-state Tau interactions that differ between wild-type Tau (TauWT) and Tau with familial FTD mutations, we next used affinity purification-mass spectrometry (AP-MS). Using the same strategy for genetically modified i^3^Neurons with TauWT (Figure 1A), we generated iPSCs with inducible expression of either TauP301L or TauV337M with an N-terminal flag tag (Figure S4A) and normal karyotypes (Figure S4B and S4C). Six weeks after the induction of neuronal differentiation, doxycycline treatment was used to turn on expression of flag-tagged TauWT, TauP301L, and TauV337M for 24 hours before the cells were lysed (Figure 5A). A flag pulldown was performed on cell lysates, followed by purification, trypsinization, and mass spectrometry analysis (Table S4). Importantly, the same levels of flag-tagged TauWT, TauP301L and TauV337M proteins were immunoprecipitated as detected with mass-spec, allowing us to compare the effects of mutations on tau interactome (Figure 5B).

**Figure 5.**
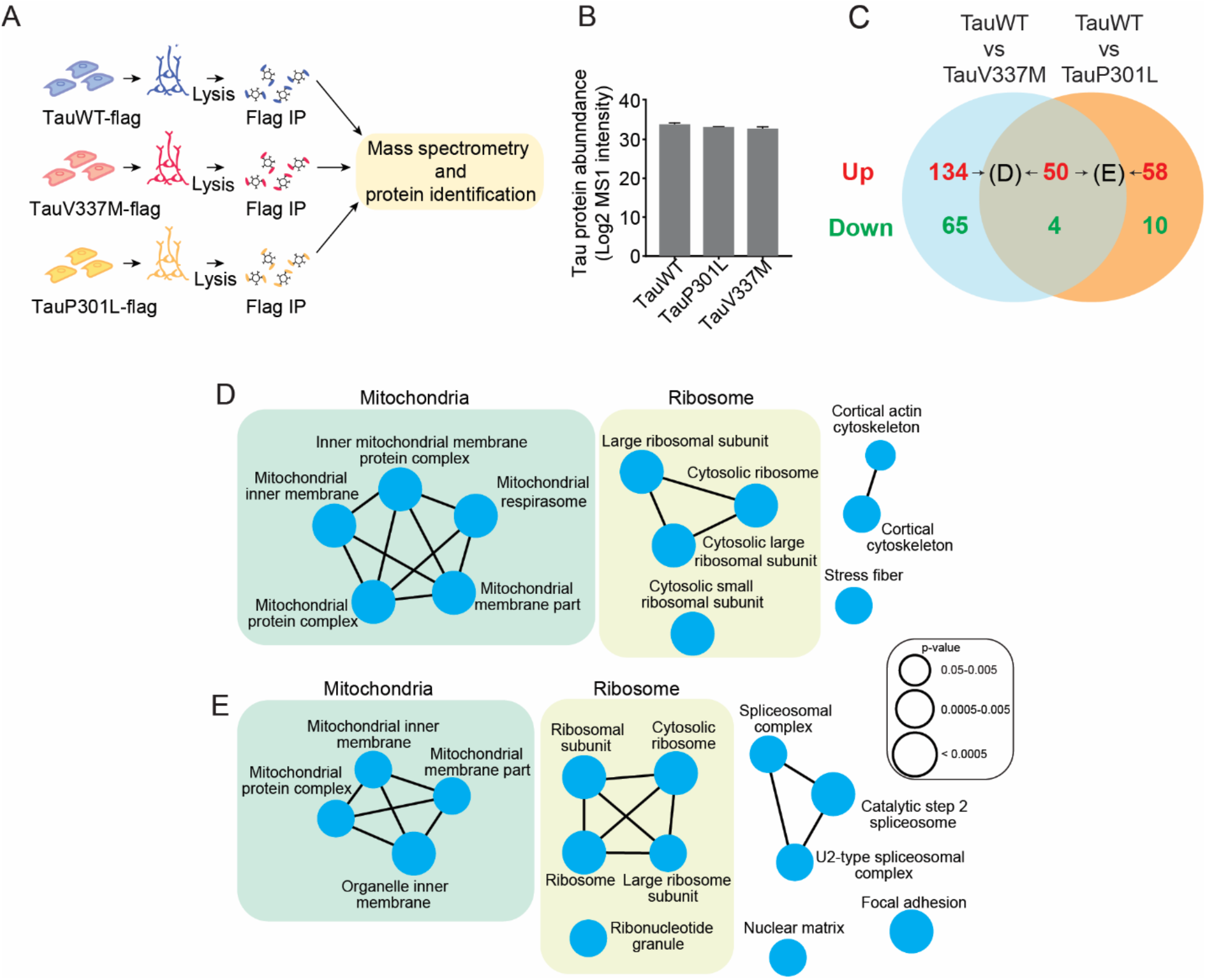
Comparison of the Tau interactome in the context of wild-type and FTD-associated mutant Tau by affinity purification mass spectrometry. **(A)** Schematic of the workflow for the affinity purification identification of Tau-associated proteins by mass spectrometry. **(B)** Expression levels of flag-tagged TauWT, TauP301L and TauV337M detected by mass spectrometry in the AP-MS experiments. See also Figure S4. **(C)** Venn diagram indicating the number of proteins identified as differential interactors between TauWT and mutant tau (TauV337M and TauP301L). In red, proteins that preferentially interact with tauWT. In green, proteins that preferentially interact with mutant tau. See also Table S5. **(D)** ClueGO cellular compartment pathway enrichment of the 184 proteins found to preferentially interact with TauWT with respect to TauV337M (labeled as “D” in panel C). **(E)** ClueGO cellular compartment pathway enrichment of the 108 proteins found to preferentially interact with TauWT with respect to TauP301L (labeled as “E” in panel C).

We focused our analysis of the AP-MS interactomes on those interactions that were lost or gained with an FTD mutation on Tau, either TauV337M or TauP301L (Figure 5C). Notably, the V337M and P301L mutations led to loss of interaction with 184 and 108 proteins, respectively, as compared to TauWT (Figure S5). Both mutations resulted in lost interacting proteins including ribosomal and mitochondrial proteins (Figures 5D and 5E), suggesting a converging disease mechanism involving a difference in the interactome of mutant pathogenic tau that could affect neuronal function.

The mitochondrial proteins that were more strongly associated with TauWT than with the Tau mutants are mostly located within the internal mitochondrial membrane (IMM) (Figure 6A). For example, TauWT interacted stronger with distinct protein complexes critical in mitochondrial bioenergetics including several subunits of complex I, III, IV and ATPase (or complex V) of the electron transport chain (ETC) (Figure 6A). Both TauV337M and TauP301L reduced the interaction with Cytochrome C (CYCS). Other IMM mitochondria proteins also exhibited stronger interaction with TauWT than TauV337M, TauP301L or both. TauP301L exhibited weaker interactions with SLC25A4, SLC25A5 and SLC25A6, distinct Adenine Nucleotide Translocators (ANT) that transport the ATP generated within the mitochondria to the cytoplasm (Figure 6A), consistent with the finding that SLC25A4 interacts with an N-terminal truncated fragment of Tau in synaptic mitochondria of AD patients (Amadoro et al., 2012b). Compared with TauV337M, TauWT interacts more strongly with HADHA and HADHB, two components of the mitochondrial trifunctional protein that catalyzes the beta oxidation of fatty acids into acetyl-CoA (Figure 6A). TauWT also showed increased interaction with several amino acid transporters as compared to TauV337M, including SLC25A13 and MPC2, which control the mitochondrial levels of glutamate and pyruvate, respectively (Figure 6A). Pyruvate is another precursor of acetyl-CoA, the starting molecule of the tricarboxylic acid (TCA) cycle, further supporting a role of Tau in mitochondria metabolism.

**Figure 6.**
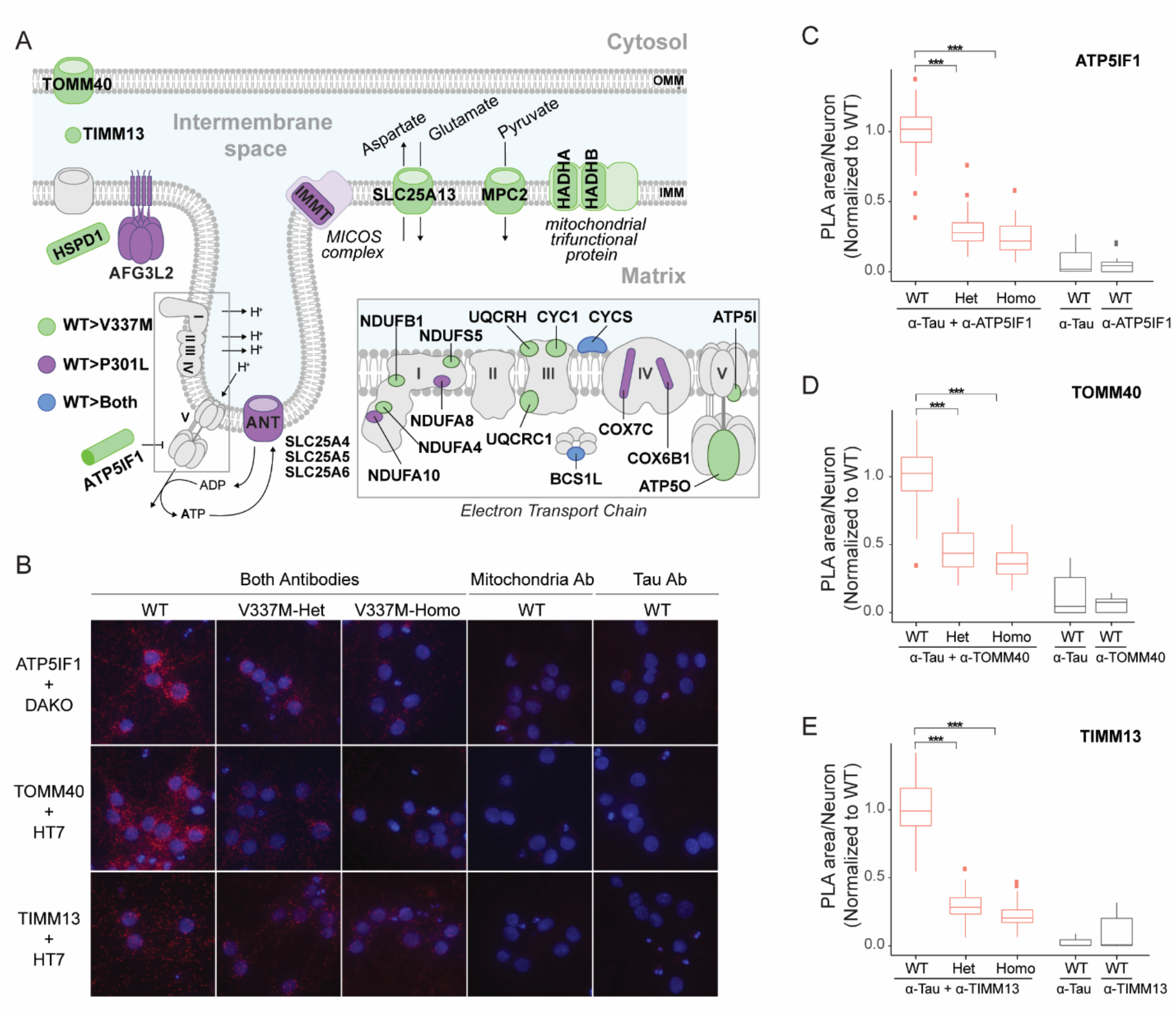
Validation of the mitochondrial localization of TauWT in human neurons. **(A)** Illustration of the TauWT-preferential interactors that locate in the mitochondrial membrane. In green, proteins found to preferentially interact with TauWT with respect to TauV337M. In purple, proteins found to preferentially interact with TauWT with respect to TauP301L. In blue, proteins found to preferentially interact with TauWT with respect to both TauV337M and TauP301L. OMM=Outer mitochondrial membrane, IMM= Inner mitochondrial membrane. **(B)** Representative images of PLA reactions between ATP5IF1, TOMM40 or TIMM13 and TauWT, TauV337M-Het or TauV337M-Homo (detected with either Dako or HT7 antibodies) in 5-week old human iPSC-derived neurons. PLA reaction controls only contain either the anti-mitochondria or anti-Tau antibodies. PLA signal=red, DAPI=blue. **(C-E)** Quantification of the area of the PLA reaction per neuron between ATP5IF1 and Tau **(C)**, TOMM40 and Tau **(D)** and TIMM13 and Tau **(E)** from the experiments depicted in B. Values from 3 independent experiments with at least 150 neurons/group analyzed in each experiment from 2 technical replicates are represented as boxplots. Tukey’s *post*-*hoc* analyses were used to compare statistical differences between designated groups after fitting a linear mixed-effects model, *** p < 0.001.

We next validated the interaction of TauWT with mitochondria proteins and the effects of V337M mutation, which exerts a stronger effect on TauWT interactome than P301L. We performed PLA between Tau and distinct mitochondrial proteins using isogenic human iPSC-derived neurons carrying WT, V337M-Heterozygous (V337M-Het) or V337M-Homozygous (V337M-Homo) mutations at the endogenous *MAPT* locus (Sohn et al., 2019; Wang et al., 2017a). Three mitochondrial proteins from the AP-MS experiment were selected based on their localizations: TOMM40, located at the Outer Mitochondrial Membrane (OMM) where it regulates the import of proteins into mitochondria, TIMM13, located in the Intermembrane Space (IMS) where it serves as a chaperone for imported proteins, and ATP5IF1, which localizes to the IMM and constitutes an inhibitor of the ETC ATPase (Figure 6A). The interaction between TauWT and the selected mitochondrial proteins was confirmed with the PLA experiments (Figure 6B-E). Negative controls including only the antibody to detect Tau or the antibodies to detect the mitochondrial proteins confirmed the specificity of the PLA signal (Figure 6B-E). Quantification of PLA signal showed that TauWT exhibited stronger interactions with TOMM40 (figure 6C), TIMM13 (Figure 6D), and ATP5IF1 (Figure 6E), than TauV337M. Together, our results provide direct evidence that TauWT’s interaction with mitochondria proteins is weakened by FTD mutations.

### TauV337M neurons have altered mitochondrial bioenergetics

To assess if the altered Tau interactions with mitochondria proteins are associated with functional consequences, we next evaluated mitochondrial bioenergetics in isogenic human iPSC-derived neurons carrying WT, V337M-Het or V337M-Homo mutations. Since V337M mutation weakened interactions with mitochondria proteins along ETC, we measured the mitochondrial membrane potential (ΔΨm)-dependent TMRM accumulation in the mitochondria of 4-week-old neurons (Figure 7A, B). V337M-Het and V337M-Homo neurons showed reduced ΔΨm compared to WT neurons (Figure 7C), suggesting either a decrease of the activity of complexes I-IV of the ETC, or an increased proton flux across the IMM of mutant tau neuronal mitochondria.

**Figure 7.**
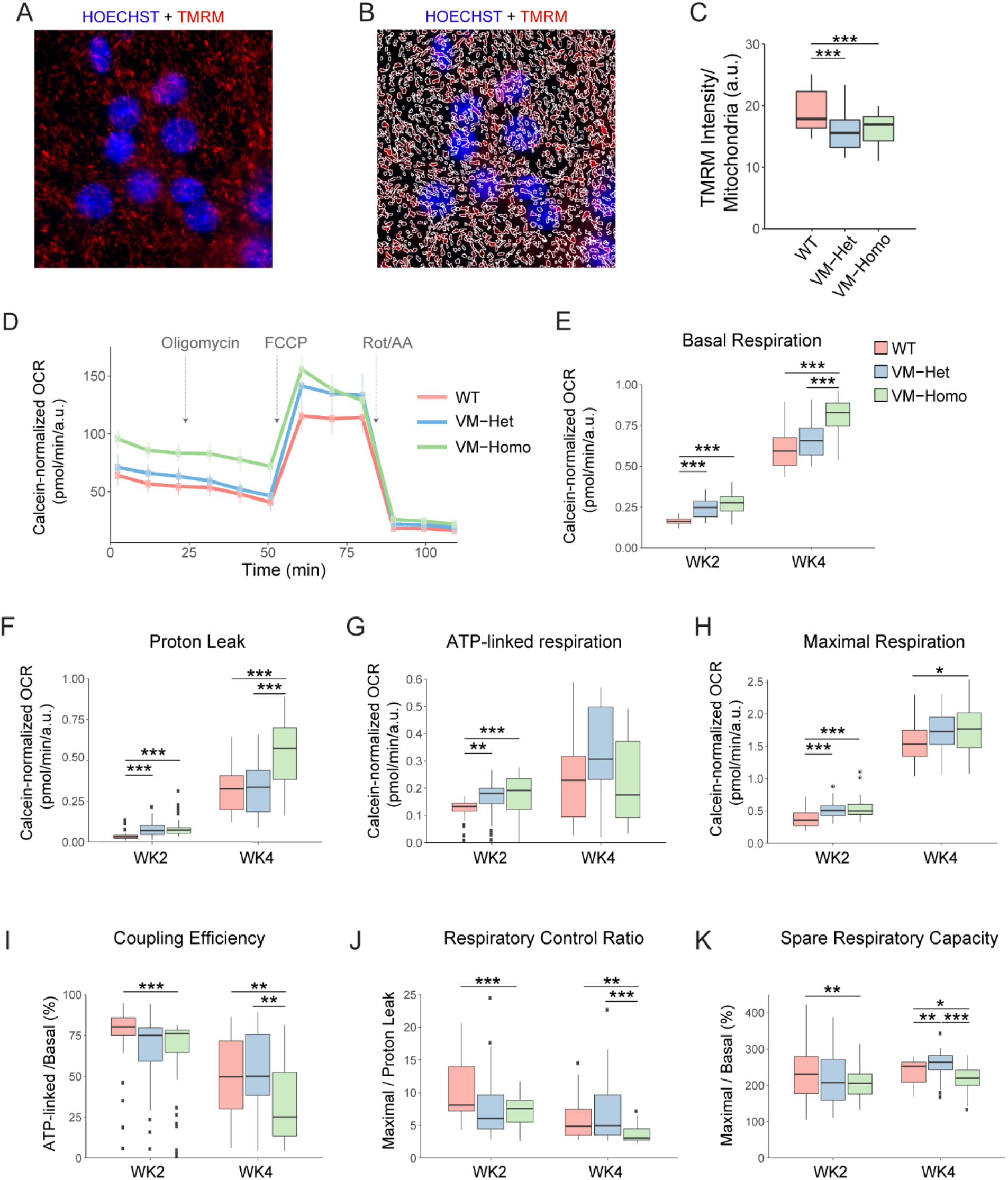
TauV337M neurons display mitochondrial bioenergetic alterations. **(A)** Representative image of 4 week-old human neurons incubated with TMRM (red) and Hoechst 33342 (blue). **(B)** Example of TMRM masking for intensity analysis of the image represented in (A). **(C)** TMRM intensity analysis of TauWT (WT), TauV337M-Het (VM-Het) and TauV337M-Homo (VM-Homo) 4 week-old neurons under baseline conditions or treated with oligomycin. 2 independent experiments with at least 8 internal replicates for each genotype and treatment were used. At least 30,000 mitochondria were analyzed per internal replicate. Tukey’s *post*-*hoc* analyses were used to compare statistical differences between designated groups after fitting a linear mixed-effects model, *** p < 0.001. **(D)** Representative OCR measurements obtained from Seahorse assays with TauWT (WT), TauV337M-Het (VM-Het) and TauV337M-Homo (VM-Homo) 4 week-old neurons normalized to calcein intensity. Arrows indicate addition of oligomycin, FCCP and Rotenone+Antimycin A (Rot/AA). **(E-F)** Quantification of basal respiration **(E)**, ATP-linked respiration **(F)**, maximal respiration **(H)** and proton leak **(F)** from Seahorse experiments depicted in (D) with 2- and 4-week-old neurons. At least 3 independent experiments with 8 technical replicates were conducted per condition. Tukey’s *post*-*hoc* analyses were used to compare statistical differences between designated groups after fitting a linear mixed-effects model, *** p < 0.001, ** p<0.01, *p<0.5. **(I-K)** Coupling efficiency **(I)**, respiratory control ratio **(J)**, and spare respiratory capacity **(K)** were calculated using the indicated parameters from the Seahorse experiments depicted in D-F. Tukey’s *post*-*hoc* analyses were used to compare statistical differences between designated groups after fitting a linear mixed-effects model, *** p < 0.001, ** p<0.01, *p<0.5.

To further dissect the effects of V337M mutation on the functionality of the ETC, we next measured the oxygen consumption rate (OCR) in two and four week-old neurons upon exposure to different stressors (Figure 7D) (Brand and Nicholls, 2011). A dramatic increase in basal (Figure 7E) and maximal respiration (Figure 7H) was observed when comparing four week-old with two-week-old neurons, consistent with a metabolic shift from aerobic glycolysis to oxidative phosphorylation and/or an increase in mitochondria number associated with neuronal differentiation and maturation (Zheng et al., 2016). V337M-Homo neurons showed higher levels of basal respiration at both 2- and 4-weeks post differentiation compared to WT neurons (Figure 7E). While ATP-linked respiration was not significantly different amongst genotypes in 4-week-old neurons (Figure 7G), V337M-Homo neurons showed significantly elevated proton leak not associated with energy generation, supporting elevated uncoupling of V337M-Homo mitochondria (Figure 7F). Upon depolarization of the IMM with the mitochondrial uncoupler FCCP, V337M-Homo neurons showed a small, but significant increase in the maximal respiration compared to WT neurons (Figure 7H), suggesting that the ETC from V337M-Homo mitochondria can work as efficiently as that in WT mitochondria in conditions that mimics an acute energy demand.

To gain more insight into how the V337M mutation could impact mitochondria energy supply, we next analyzed coupling efficiency, respiratory control ratio, and spare respiratory capacity, three internally normalized parameters which are more sensitive to mitochondria dysfunction (Brand and Nicholls, 2011; Connolly et al., 2018; Divakaruni et al., 2014). V337M-Homo mitochondria used a higher fraction of the basal respiration to oxidize substrates not destined to drive ATP synthesis, as indicated by the diminished coupling efficiency (Figure 7I). We also found a significant reduction in the respiratory control ratio of V337M-Homo mitochondria (Figure 7J), which is sensitive to changes in substrate oxidation and proton leak but independent of ATP synthesis (Brand and Nicholls, 2011; Connolly et al., 2018; Divakaruni et al., 2014). Moreover, the spare respiratory capacity, which indicates how close a cell is to its bioenergetic limit, was significantly reduced in V337M-Homo mitochondria compared with WT mitochondria (Figure 7K). Altogether, these functional analyses showed that the reduced mitochondria interactions caused by the V337M mutation could compromise the neuron’s ability to sustain prolonged energy demand, supporting the notion that the TauWT interaction with mitochondria proteins may play an important physiological function.

### Levels of Tau interactors modified by FTD mutation correlate with disease severity in human AD brain

Recent work identified six co-expressed protein modules that significantly associate with Alzheimer’s disease progression using a multi-omics approach on multiple cohorts from the Accelerating Medicines Partnership – Alzheimer’s Disease (AMP-AD) consortium (Swarup et al., 2020). These protein co-expression modules are conserved across multiple data sets, including FTD and PSP and in controls, and thus represent robust biological processes in brain that are differentially regulated in dementia involving Tau. We leveraged this unbiased, genome-wide data set to probe the pathophysiological relevance of the proteins that show reduced interaction with FTD mutant Tau in human brain. Interestingly, 29 out of the 108 proteins (Methods, permutation test p=0.009) and 48 out of the 184 proteins (Methods, permutation test p=0.0017), which preferentially interact with TauWT compared to TauP301L and TauV337M, respectively, were represented in the modules associated with AD.

Analyses of the proteomic data from the Banner Sun Health Research Institute (Banner) cohort revealed that there is a significant reduction in the levels of TauWT-preferential interactors in AD patients compared to control, asymptomatic, mild-cognitive impairment (MCI) patients (Figure 8A, TauWT>TauV337M AD vs control p=4×10^-15^, TauWT>TauP301L AD vs control p=4×10^-11^). Moreover, the levels of these Tau interacting proteins were negatively correlated with two distinct neuropathological scores: the Consortium to Establish a Registry for Alzheimer’s Disease (CERAD) score (Figure 8B), and the Braak score (Figure 8C). Similar correlations were observed in the Baltimore Longitudinal Study of Aging (BLSA) cohort (Figure. S6A-C, TauWT>TauV337M AD vs control p=5×10^-5^, TauWT>TauP301L AD vs control p=6×10^-4^). Altogether, this data suggests that proteins that normally interact with TauWT whose interaction is reduced upon mutations in Tau may play an important role in AD pathogenesis, where reduced levels correlate with disease severity. They also show the relevance of these findings to changes occurring in vivo in human brain.

**Figure 8.**
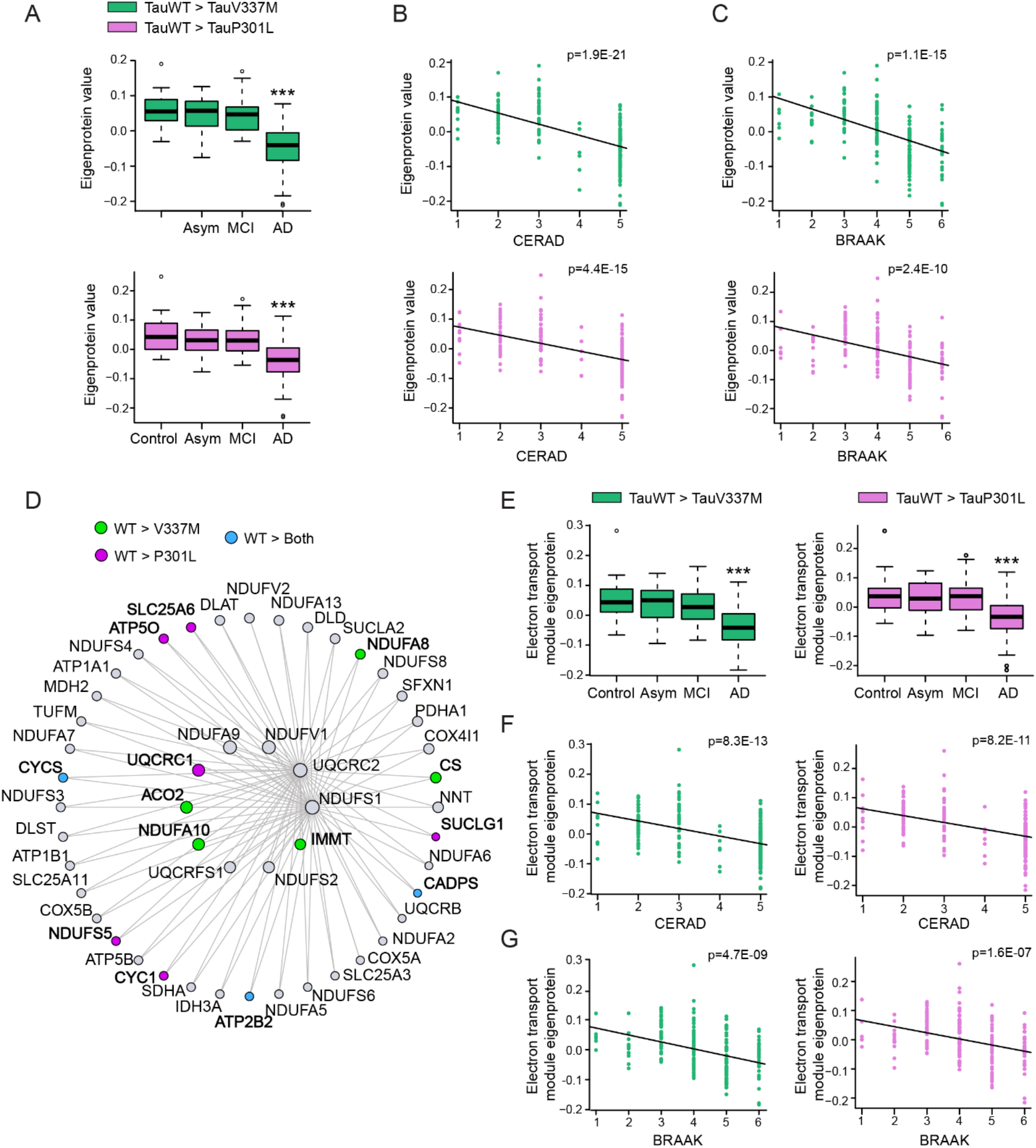
Decreased expression of TauWT-preferential interactors correlates with human AD progression. **(A-C)** Plots showing TauWT-preferential interactor eigenprotein trajectory with AD diagnosis **(A)**, CERAD **(B)**, and BRAAK **(C)** scores in Banner cohort. TauWT-preferential interactors as compared to TauV337M and P301L are colored as green and pink, respectively. ***p < 0.005. See also Figure S6. **(D)** Co-expression graph of the first 50 hub proteins of the C2-mitochondrial module showing TauWT-preferential interactors as compared to TauV337M (green), P301L (pink) or both mutants (blue). **(E-G)** Plots showing C2-mitochondrial module TauWT-preferential interactor eigenprotein trajectory with AD diagnosis **(E)**, CERAD **(F)**, and BRAAK **(G)** scores in Banner cohort. TauWT-preferential interactors as compared to TauV337M and P301L are colored as green and pink, respectively. ***p < 0.005. See also Figure S6.

Given the abundance of mitochondrial proteins that were TauWT-preferential interactors, we further explored the dementia associated AMP-AD neuronal-specific C2 module, which is enriched for mitochondrial electron transport chain subunits (Swarup et al., 2020). We found that 25 and 13 TauWT-preferential interactors, compared to TauV337M and TauP301L, respectively, are components of the C2 module (Figure 8D). The total protein levels of these C2-TauWT-mitochondrial interactors were also significantly reduced in AD patients compared to control, asymptomatic, MCI patients (Figure 8E, TauWT>TauV337M-C2 AD vs control p=1×10^-9^, TauWT>TauP301L-C2 AD vs control p=8×10^-8^). Comparison with clinical progression revealed negative correlations between the levels of C2-TauWT-mitochondrial interactors and CERAD and Braak neuropathological scores (Figure 8F and 8G). Analysis of the levels of C2-TauWT-mitochondrial interactors in the Baltimore Longitudinal Study of Aging (BLSA) cohort showed a comparable effect (Figure S6D-F, TauWT>TauV337M-C2 AD vs control p=0.03, TauWT>TauP301-L-C2 AD vs control p=0.01). Altogether, these data suggest that the C2-TauWT-mitochondrial interactors identified *in vitro* using human neurons are downregulated during AD progression and are associated with both cognitive decline and the accumulation of pathological Tau in the brain.

## DISCUSSION

Our proteomic analyses reveal the multi-modal and dynamic Tau interactome with unprecedented spatiotemporal resolution in human neurons. APEX proximity-dependent labeling with mass spectrometry enabled the identification of specific interactions in the Tau interactome involving either the N-or the C-terminal domain of Tau at the amino acid level. Enhanced neuronal activity rapidly changed the Tau interactome, causing Tau to become more associated with vesicle and SNARE complex-associated proteins during the activity-induced secretion of Tau from neurons. Analyses of the Tau interactome with or without FTD-causing mutations revealed that TauWT has a greater association with mitochondrial proteins than FTD mutant Tau. We found that these TauWT-interacting mitochondrial proteins are components of a key protein module that is downregulated in human AD brains, and that human neurons expressing the same levels of pathogenic Tau have abnormal mitochondrial bioenergetics. From complementary mass spectrometry approaches to map subcellular, activity-induced, and mutation-dependent differences in the Tau interactome, our findings provide novel insight into the role of Tau and Tau-interacting proteins in disease.

APEX-mediated labeling of Tau-interacting proteins was performed in living neurons with intact cellular compartments maintaining the endogenous subcellular distribution of the Tau interactome. Thus, Tau protein-protein interactions that occur more strongly within intact compartments or that bind transiently could be preferentially detected in a snapshot of time. Detection of numerous presynaptic vesicle-associated proteins at the active zone including Syntaxin-1A/B, SNAP25, Munc18, Mint1 and α-Synuclein (Figure 3E), and Synaptotagmin-1 which was biotinylated following enhanced neuronal activity (Figure 4F), suggests that Tau is closely associated with docked vesicles at sites of vesicle fusion in the presynaptic terminal. Consistent with studies showing that Tau can bind directly to synaptic vesicles (McInnes et al., 2018; Zhou et al., 2017), we identified biotinylation on 5 integral synaptic vesicle proteins that span the vesicle membrane including SV2A, VGLUT2, Synaptogyrin-1, Synaptotagmin-1 and SV2C. Importantly, the tyrosines biotinylated by APEX Tau on these proteins were all located on the cytosolic domain of the proteins rather than the lumenal domains supporting that the Tau protein-protein interaction occurred on the cytosolic side of the vesicle. The interaction of the N-terminal of Tau with Synaptogyrin-3 has been reported to inhibit presynaptic vesicle release (McInnes et al., 2018). We did not detect Synaptogyrin-3 in our Tau interactome, however, Synaptogyrin-1, which is found on the same vesicles as Synaptogyrin-3 (Belizaire et al., 2004) was biotinylated specifically by N-APEX Tau. Previous studies also showed that Tau interacts with RNA binding proteins (Maziuk et al., 2018) and forms droplets with RNA (Zhang et al., 2017). In our study, APEX-Tau was found in proximity to RNA granule proteins including G3BP2, hnRNPA1, PABPC1, and CIRBP as well as the chaperone complex proteins HSP70 and HSP90, which are known Tau binding proteins.

APEX proximity dependent mapping revealed distinct protein-protein interactions with the N- and C-terminal domains of Tau which may be mediated in part by Tau fragments, or by domain specific interactions. It is estimated that APEX2 biotinylates proteins up to 10-20 nm away (Hung et al., 2014; Rhee et al., 2013), whereas the length of the Tau protein is approximately 63-70 nm (Hagestedt et al., 1989; Ruben et al., 1991), thus allowing for the spatial resolution of interactions in proximity to the N- or C-terminus. N-terminal Tau enriched interactions included proteins that regulate post-synaptic organization, dendritic spines, vesicle transport and proteasome-dependent protein degradation. C-terminal enriched interactions included proteins that mediate vesicle fusion at presynaptic sites and the cellular stress response. The domain-specific interactions detected here are different from those reported in a proteomics study on fragments of the N-terminal and C-terminal domains of Tau expressed in SH-SY5Y cells (Gunawardana et al., 2015), possibly because the neuroblastoma cells do not have the physiological and morphological features of mature human neurons.

Our findings revealed the dynamic protein-protein interactions with Tau during activity-induced Tau secretion. We confirmed that neuronal network activity drives Tau secretion from neurons (Wu et al., 2016; Yamada et al., 2014), and that Tau secretion is calcium-dependent (Pooler et al., 2013). Most of the Tau protein interactions remained intact in the neurons following enhanced activity, consistent with the majority of Tau remaining inside the cell. However, we discovered that neuronal activity enhanced the interaction of Tau with the SNARE complex-associated proteins Synaptotagmin-1 and Mint1 and the synaptic vesicle proteins SV2C, Synapsin 1, and RAB3A suggesting that that presynaptic vesicle fusion machinery could regulate activity-dependent Tau release through a direct protein-protein interaction. Depolarization of presynaptic terminals causes a local influx of calcium that binds to Synaptotagmin-1 on synaptic vesicles docked at the active zone to trigger vesicle release. Synaptotagmin-1 interacts with Syntaxin1/SNAP25 in the SNARE complex which when triggered forces the plasma membrane to break for vesicle fusion. Activity-induced Tau release from neurons is inhibited by tetanus toxin which disrupts the SNARE complex function (Pooler et al., 2013), suggesting that the SNARE complex plays an key role in Tau release. Our results show for the first time that Tau associates directly with Synaptotagmin-1 and the SNARE complex when it is secreted from the depolarized presynaptic terminal. Another proposed mechanism for activity-induced Tau release involves exosome-mediated release of Tau and uptake across synaptic connections (Wang et al., 2017c). In human neuron cultures the majority of secreted Tau is free-floating with a smaller amount of Tau detected in extracellular vesicles (Guix et al., 2018), indicating that there could be multiple mechanisms contributing to Tau release. To what extent Tau release at presynaptic terminals involves vesicular or vesicle-free routes remains to be determined.

Our AP-MS results identified a broader set protein-protein interaction networks for Tau in human neurons. Previous mass spectrometry studies used AP-MS in Tau transgenic mice and Tau-overexpressing cell lines to reveal RNA-binding proteins as Tau interactors (Maziuk et al., 2018), and to shed light on specific regulators of ubiquitination and protein degradation that bind directly to Tau (Choi et al., 2020; Gunawardana et al., 2015; Thompson et al., 2012; Wang et al., 2017b). In accordance with those observations, we found that Tau in human neurons interacts with RNA-binding proteins (Table S4), although the interaction of TauV337M and TauP301L with some ribosomal proteins is partially reduced (Figure 5D and 5E). Mutant Tau also had a reduced interaction with ribosomes in an FTD mouse model (Maziuk et al., 2018). However, Tau showed increased affinity to ribosomes in microsomes from AD patients as compared to control patients (Meier et al., 2016). Whilst the discrepancies described above might be explained by the models and techniques used to characterize these interactions, cumulatively the data point toward an important role of Tau in ribonucleoprotein organization and ribosomal functions that might contribute to disease pathogenesis.

Our AP-MS data established extensive interactions of Tau with proteins involved in mitochondrial bioenergetics, including many subunits of the ETC and SLC25A4, a transporter that controls the exchange of cytoplasmic ADP with mitochondrial ATP across the IMM, which was previously described to interact with Tau in synaptic mitochondria of AD patients (Amadoro et al., 2012a). Our combined AP-MS study and PLA assays demonstrate for the first time that FTD-linked mutations diminished the interaction with mitochondria proteins at the OMM, and especially those at IMM, such as distinct subunits from Complex I, III, IV, V and Cytochrome C of the ETC. The interaction between endogenous mouse Tau and mitochondrial proteins has been reported previously (Liu et al., 2016; Wang et al., 2017b). TauWT has been shown to interact with lipid membranes (Brandt et al., 1995; Georgieva et al., 2014), and post-translational modifications associated with mutant Tau diminish this association (Ekinci and Shea, 2000; Gauthier-Kemper et al., 2011; Maas et al., 2000). It is possible that the broad interaction between IMM proteins and TauWT reflects a physiological accumulation of TauWT in the mitochondrial membranes, which is disrupted upon Tau mutations. The diminished interactions in V337M neurons were associated with alterations in mitochondria bioenergetics with decreased coupling efficiency, which could impact their ability to maintain ATP levels under prolonged energetic stress. Indeed, tauopathy mice expressing P301L mutant Tau exhibited reduced respiratory control ratio and impaired ATP synthesis with aging (David et al., 2005), and P301L-overexpressing SH-SY5Y cells exhibited decreased mitochondrial ATP levels and increased susceptibility to oxidative stress (Schulz et al., 2012).

Our study further established that levels of TauWT-preferential interactors negatively correlated with clinical and pathological disease progression in two different cohorts representing control, asymptomatic AD, MCI and AD patients (Swarup et al., 2020). Interestingly, one of the two neuronal-specific proteomic modules downregulated in neurodegeneration consist of mitochondrial proteins from the ETC (Swarup et al., 2020), and includes several of the mitochondrial TauWT-preferential interactors. Altogether, analysis of human AD brains indicates that TauWT-preferential interactors might play an important role in disease development, and it is possible that either a reduction of the levels of these proteins or their decreased association with mutant Tau have similar effects on the cellular bioenergetic homeostasis.

Despite the broad physiological consequences of pathogenic Tau in disease, development of tau-targeted therapeutic strategies has been limited by lack of understanding of how Tau directly mediates these processes. The dynamic Tau interactome reported here maps the complex multifaceted nature of the Tau protein function in disease related processes. Our work provides an extensive resource for possible mechanisms by which Tau directly influences cellular functions through protein-protein interactions in human neurons, which pave the way for novel tau-targeted strategies to counteract neurodegenerative processes.

## AUTHOR CONTRIBUTIONS

L.G., T.E.T. and J.M.P. conceived the project. L.G., T.E.T., J.M.P., and D.L.S. designed experiments. T.E.T., J.M.P., D.L.S., M.M., E.S., G.K., L.S.M., M.M., and X.C. performed experiments. T.E.T., J.M.P., T.S.C., M.E.W., R.H., E.H., S.A.M., M.T., S.W.M., C.W., P-D.S., J.M., Y.Z., D.G., D.L.S., N.J.K. developed experimental protocols, tools or reagents or analyzed data. T.E.T., J.M.P. and L.G. wrote the manuscript.

## ACKNOWLEDGEMENTS

We thank Dr. Bruce Conklin (Gladstone Institutes) for the WTC11 iPSCs, pUCM vector and TALENS, Connor Ludwig for technical support, Berenice Rossi for graphic design, Nathan Basisty for advice on analyses, and Erica Delin for administrative assistance. This work was supported by the NIH (U54NS100717 to L.G. and N.J.K., K01 AG057862 to T.E.T.), NINDS (R25NS065723 Translational Neuroscience Training Grant), Fundación Ramón Areces (postdoctoral fellowship to J.M.P) and a grant from the Tau Consortium (to L.G.).

The results published here are in whole or in part based on data obtained from the AMP-AD Knowledge Portal (https://adknowledgeportal.synapse.org/). BLSA data were generated from postmortem brain tissue collected through the National Institute on Aging’s Baltimore Longitudinal Study of Aging and provided by Dr. Levey from Emory University.

Banner data were provided by Dr. Levey from Emory University. A portion of these data were generated from samples collected through the Sun Health Research Institute Brain and Body Donation Program of Sun City, Arizona. The Brain and Body Donation Program is supported by the National Institute of Neurological Disorders and Stroke (U24 NS072026 National Brain and Tissue Resource for Parkinson¹s Disease and Related Disorders), the National Institute on Aging (P30 AG19610 Arizona Alzheimer¹s Disease Core Center), the Arizona Department of Health Services, United States (contract 211002, Arizona Alzheimer¹s Research Center), the Arizona Biomedical Research Commission, United States (contracts 4001, 0011, 05-901 and 1001 to the Arizona Parkinson’s Disease Consortium) and the Michael J. Fox Foundation for Parkinson’s Research, United States.

## EXPERIMENTAL PROCEDURES

### Generation of iPSC clones

We used human iPSCs described in a previous study (Wang et al., 2017a), that were engineered for inducible expression of Ngn2 from a transgene integrated in the AAVS1 locus of WTC11 cells with a wild-type genetic background (Miyaoka et al., 2014). The iPSC clone used contained one copy of the Ngn2 cassette integrated in a single allele of the AAVS1 locus. Cre recombinase treatment was used to excise the puromycin resistance gene from the iPSC clone carrying the Ngn2 transgene. The sequence for APEX2 with a flag tag was cloned into either the N- or C-terminus of wild-type human Tau (2N4R) as well as the N-terminus of Tubulin α 1b, and the N-termini of TauP301L and TauV337M (2N4R). The N-APEX Tau, C-APEX Tau, N-APEX TauP301L and N-APEX TauV337M sequences were each subcloned into a pUCM vector with AAVS1 homology arms and the Tet-On 3G tetracycline-inducible expression system as described (Wang et al., 2017a). The pUCM donor vector together with TALEN pairs (kind gifts from Dr. Bruce Conklin, Gladstone Institutes) were used for targeted integration of each cassette into the second allele of the AAVS1 locus. The Ngn2 integrated iPSCs were first grown in E8 media, dissociated with Accutase and then transfected with one of the Tau or Tubulin pUCM vectors along with AAVS1 TALEN pairs using a Human Stem Cell Nucleofector Kit 1 (Lonza) with the Nucleofector 2b Device (Lonza). Puromycin (0.1-0.3 ug/mL) was added 24-48 hours later to select for cells with genetic integration. E8 media with puromycin was regularly replenished until isolated and established colonies were formed. An EVOS FL microscope (Invitrogen) was used to pick colonies for individual clones. Clones were then tested for APEX-tagged Tau or Tubulin expression by immunoblotting and immunocytochemistry. The use of iPSCs in this study was approved by the Committee on Human Research at the University of California, San Francisco (15-15798).

### Cell culture

Pre-differentiation of iPSCs into neurons was initiated by plating 2 ×10^6^ iPSCs in each well of matrigel-coated 6-well plate with Knockout DMEM/F-12 media containing doxycycline (2 ug/mL), N2 supplement, non-essential amino acids, brain-derived neurotrophic factor (10 ng/mL, Peprotech), neurotrophin-3 (10 ng/mL, Peprotech) and ROCK inhibitor (Y-27632, Cayman chemicals). The media was replaced the next day without ROCK inhibitor, and pre-differentiation was maintained for a total of three days. On day 0 the predifferentiated precursor cells were dissociated with Accutase and plated onto matrigel coated coverslips or tissue culture plates for the growth of neuron cultures in Neurobasal-A media containing B27 supplement, Glutamax, BDNF (10 ng/mL) and NT3 (10 ng/mL) with doxycycline (2ug/mL). For proteomics experiments, 8 x 10^6^ precursor cells were plated in a 10 cm tissue culture plate for each replicate. Rat astrocytes were added to the neurons on day 1, and half of the media was replaced on day 5 with supplemented Neurobasal-A media containing cytosine β-D-arabinofuranoside (Ara-C) but lacking doxycycline. On day 10, half of the media was removed and three times the remaining volume in the culture was replenished with MEM media containing glucose (27.7mM), NaHCO_3_ (2.4mM), B-27 supplement, L-glutamine, Ara-C) and fetal bovine serum (5%). One third of the supplemented MEM media was replaced every week thereafter. At 5-7 weeks of age, doxycylcine (2 ug/mL) was added to the neurons 24 hours before experiments were performed to express the APEX2-flag tagged Tau or Tubulin proteins. For experiments performed to assess mitochondrial function, human iPSC-derived neurons were differentiated from wild-type (WTC11) iPSCs carrying the NGN2 transgene and isogenic iPSCs that were edited by CRISPR/Cas9 to generate heterozygous or homozygous V337M mutations as previously described (Sohn et al., 2019).

### Western blot

Following APEX stimulation and quenching, human iPSC-derived neuron cultures were washed with cold PBS, then homogenized in cold RIPA buffer containing 50 mM Tris, pH 7.5, 150 mM NaCl, 0.5% Nonidet P-40, 1mM EDTA, 1 mM phenylmethyl sulfonyl fluoride, protease inhibitor cocktail, phosphatase inhibitor cocktail. Lysates were sonicated, centrifuged at 18,000 g at 4°C for 15 min, and the supernatant was collected. Protein concentration was determined by Bradford assay (Bio-Rad). Equal amounts of protein for each sample were loaded and run on a 4–12% SDS-PAGE gel (Invitrogen). A nitrocellulose membrane (GE Healthcare) was used for transfer of proteins, and blots were blocked with 5% milk, and incubated with primary antibodies for mouse anti-flag (Sigma), HT7 (Thermo Fisher), GAPDH (Millipore) and streptavidin horseradish peroxidase conjugated (HRP, Thermo Fisher). Secondary HRP antibodies (Millipore) and chemiluminescence (Pierce) were used for detection of immunoblotting.

### APEX proximity-dependent labeling and enrichment of biotinylated proteins

To start proximity-dependent labeling by APEX, the neuronal media was removed and cells were washed with extracellular solution warmed to 37°C containing (in mM): 140 NaCl, 5 KCl, 2.5 CaCl_2_, 2 MgCl_2_, 10 HEPES, 10 glucose at pH 7.4. Then neurons were incubated in extracellular solution with biotin-phenol (500 µM, Adipogen) in a 37°C cell culture incubator for 30 minutes. The neurons were next treated with 1 mM hydrogen peroxide (H_2_O_2_) for 1 minute and quickly washed once with Dulbecco’s phosphate buffered saline (DPBS) followed by two washes with quenching solution consisting of 1 x PBS with 10 mM sodium azide, 10 mM sodium ascorbate, and 5 mM Trolox. The quenching solution was removed, and the neurons were either fixed for immunocytochemistry or lysed to extract proteins. The neurons were lysed at 4°C in buffer containing 50mM Tris-HCl pH 7.4, 500 mM NaCl, 0.2% SDS, 2% Triton, 1 mM DTT, a protease inhibitor tablet (Pierce), phosphatase inhibitor cocktails (Sigma), 10 mM sodium azide, 10 mM sodium ascorbate, and 5 mM Trolox. The samples were sonicated and then centrifuged at 16,500 g for 10 minutes at 4°C. The collected supernatant was then subjected to dialysis to remove excess free biotin using a Slide-A-Lyzer MINI dialysis device (Thermo Scientific). Proteins were precipitated by adding 9 parts methanol to 1 part lysate. The supernatant was discarded, and the precipitated protein pellet was washed once with methanol and then dried by vacuum centrifugation. Protein was resuspended in 8M urea, 100 mM ammonium bicarbonate, then sonicated with a probe sonicator for 1 minute to re-solubilize the proteins. A Bradford Assay was performed to determine protein concentration and 1 mg of protein was taken out to be processed further. TCEP was added to 4 mM and incubated for half an hour at room temperature. Each sample was treated with 10 mM iodoacetamide, then excess iodoacetamide was quenched with 10 mM DTT. The samples were diluted 4-fold with 100 mM ammonium bicarbonate. Trypsin (Promega) was added in a 1:100 enzyme:substrate ratio and samples were incubated overnight at 37°C with agitation. The digested samples were desalted on a 50 mg SepPak column and eluted peptides were dried thoroughly via vacuum centrifugation. Peptides were dissolved in IAP buffer containing 50 mM MOPS, 10 mM HNa_2_PO_4_, 50 mM NaCl at pH 7.5. Anti-biotin beads (Immune Chem Pharmaceuticals) were washed twice in IAP buffer before the samples were added to the beads for incubation on a rotator for two hours at 4°C. The beads were washed twice with IAP buffer and twice with H_2_O (HPLC-grade). The biotinylated peptides were eluted from the beads by 0.15% TFA with vortexing followed by a 10 minute incubation. The elution was repeated twice and the supernatants were collected for desalting on NEST C18 tips then drying by vacuum centrifugation.

### Flag pulldown of Tau

Neuron cultures were washed three times with DPBS before cold lysis buffer was added containing 50mM Tris-HCl pH 7.4, 150 mM NaCl, 1 mM EDTA, 0.5% NP40, a protease inhibitor tablet (Pierce), and phosphatase inhibitor cocktails (Sigma) at pH 7.4. The samples were lysed by 2 freeze/thaw cycles and then centrifuged at 16,500 g for 15 minutes at 4°C. Magnetic anti-flag beads (Sigma) were washed twice with IP buffer containing 50mM Tris-HCl pH 7.4, 150 mM NaCl, 1 mM EDTA (pH 7.4). Lysates were added to the beads and incubated with rotation at 4°C for 2 hours. The beads were then washed twice times with 50mM Tris-HCl pH 7.4, 150 mM NaCl, 1 mM EDTA, 0.05% NP40 (pH 7.4) and then with two additional washes using IP buffer. To elute proteins from the beads a 3xFLAG peptide (100 ug/mL, Sigma) with 0.05% RapiGest in IP buffer was added to the beads for 15 minutes at room temperature with agitation. The elution was repeated a second time and the supernatants were combined with the addition of 1.7M urea, 50 mM Tris, and 1 mM DTT. The samples were incubated at 60°C for 15 minutes, then 3mM iodoacetamide was added for 45 minutes at room temperature. After adding 3mM DTT the samples were digested with trypsin (Promega) overnight at 37°C. Trypsinized samples were acidified with 0.5% TFA then desalted on a C18 stage tip and dried by vacuum centrifugation.

### Mass spectrometry analyses

Samples were resuspended in 4% formic acid, 4% acetonitrile solution, separated by a reversed-phase gradient over a nanoflow column (360 µm O.D. x 75 µm I.D.) packed with 25 cm of 1.8 µm Reprosil C18 particles with (Dr. Maisch). Mobile phase A consisted of 0.1% FA and mobile phase B consisted of 80% ACN/0.1% FA. Biotinylated peptides were separated by an organic gradient from 12% to 22% mobile phase B over 35 minutes followed by an increase to 40% B over 40 minutes, then held at 100% B for 6 minutes at a flow rate of 300 nL/minute. Eluting peptides were directly injected into an Orbitrap Fusion Lumos Tribrid Mass Spectrometer (Thermo). Data was collected in positive ion mode with MS1 detection in profile mode in the orbitrap using 120,000 resolution, 350-1400 m/z scan range, 100 ms maximum injection, and an AGC target of 1e6. MS2 fragmentation was performed on charge states from 2-6 with a 30s dynamic exclusion, and all MS2 data was collected in centroid mode at a rapid scan rate in the ion trap with HCD (32% normalized collision energy), 15ms maximum injection time, 2e4 AGC.

Peptides resulting from FLAG-purified samples were separated by an organic gradient from 7% to 36% mobile phase B over 53 minutes followed by an increase to 95% B over 7 minutes, then held at 90% B for 6 minutes at a flow rate of 300 nL/minute. Eluting peptides were directly injected into a Q-Exactive Plus mass spectrometer (Thermo). Data was collected in positive ion mode with MS1 detection in profile mode in the orbitrap using 70,000 resolution, 350-1500 m/z scan range, 250 ms maximum injection, and an AGC target of 1e6. MS2 fragmentation was performed on charge states from 2-6 with automatic dynamic exclusion, and all MS2 data was collected in centroid mode in the orbitrap at 17,500 resolution with HCD (26% normalized collision energy), 60ms maximum injection time, 5e4 AGC.

All raw MS data were searched with MaxQuant against the human proteome (Uniprot canonical protein sequences downloaded March 21, 2018 for anti-biotin samples, or July 21, 2017 for FLAG-purified samples). Peptides, proteins, and PTMs were filtered to 1% false discovery rate in MaxQuant (Cox and Mann, 2008). MaxQuant default parameters were used with the exception that label-free quantification was turned on with match between runs set to 1 min, and for anti-biotin purified samples a variable mass addition of biotin-phenol (361.14601 Da) on Y residues was considered. Mass spectrometry data files (raw and search results) have been deposited to the ProteomeXchange Consortium (http://proteomecentral.proteomexchange.org) via the PRIDE partner repository with dataset identifier (Vizcaíno et al., 2016).

Biotinylated proteins identified in APEX experiments that were detected in fewer than half of the neuronal culture replicates (<4/7 replicates for N-APEX Tau unstimulated or 50mM KCl groups, <5/9 replicates for C-APEX Tau unstimulated or 50mM KCl groups, or <6/12 replicates for APEX-α tubulin) were removed from analyses (Tables S1 and S3). To classify Tau interactors identified by AP-MS (Table S4), proteins detected in fewer than half (<4/7 for TauWT and <4/8 for TauV337M and TauP301L) of the neuronal culture replicates for each group were removed to reduce non-specific binders. To classify proteins as TauWT, TauV337M or TauP301L preferential interactors (Table S5), proteins needed to meet one of the two following criteria: 1) To be identified in more than 50% of the replicates in both groups of the comparison and to be significantly different (adjusted P value <0.05) with a detection fold change bigger than 1.5 between genotypes, or 2) to be consistently detected in one group (more than 50% of replicates) and in 3 times more replicates than in the other genotype in the comparison (i.e. 75% of replicates in genotype A and 25% replicates of genotype B)

### Enrichment and network analyses

Gene Ontology (GO) term enrichment analyses were performed using either Gene Set Enrichment Analysis (GSEA) (Mootha et al., 2003; Subramanian et al., 2005) or ClueGO version 2.5.4 in Cytoscape version 3.7.1 (Bindea et al., 2009; Shannon et al., 2003). Enrichment and network analyses were applied from three ontologies including GO Molecular Function, GO Biological Process, and GO Cellular Compartments. The statistics used in ClueGO analyses revealed enriched networks with p-values < 0.05 from right-sided hypergeometric testing with Bonferroni correction.

### Proximity ligation assay

Human iPSC-derived neurons were grown on glass coverslips and fixed at 5-7 weeks of age in 4% paraformaldehyde in PBS for 15 min then washed three times for 5 min with PBS. The DuoLink In Situ Fluorescence (Sigma) protocol was used to perform the proximity ligation assay (PLA) on the neurons. The cultures were incubated in a humidified chamber with Duolink Blocking Solution for 1 hour at 37°C, then the blocking solution was removed and replace with the primary antibodies diluted in Duolink Antibody Diluent for 1 hour at room temperature. The primary antibodies used for PLA included Dako-Tau (Agilent Technologies), HT7 (Thermo Fisher), Munc18 (BD Biosciences), Synaptotagmin-1 (Synaptic Systems), ATP5IF1 (ThermoFisher, Cat# A-21355), TOMM40 (ProteinTech, Cat# 18409-1-AP), TIMM13 (ThermoFisher, Cat# PA5-61856).

The coverslips were washed twice for 5 min with Wash Buffer A (Sigma) then the Duolink In Situ PLA Probe Anti-Rabbit PLUS and Duolink In Situ PLA Probe Anti-Mouse MINUS PLA probes (Sigma) diluted in Duolink Antibody Diluent were applied. The coverslips were incubated in a humidified chamber with PLA probe solution for 1 hour at 37°C. Coverslips were washed twice with Wash Buffer A followed by the addition of the ligation solution for 30 min at 37°C, then another two washing steps, and incubation in amplification solution for 100 min at 37°C. The coverslips were washed twice for 10 minutes in Wash Buffer B (Sigma) and mounted onto slides with ProLong Gold Mountant (Thermo Fisher).

### Human Tau ELISA assay

Human iPSC-derived neurons were differentiated from wild-type (WTC11) NGN2 iPSCs and matured for six weeks in culture. The cell culture media was removed and replaced with extracellular solution containing (in mM): 115 NaCl, 3.5 KCl, 10 HEPES, 1 MgCl_2_, 2.5 CaCl_2_ at pH 7.4 or with high KCl extracellular solution containing (in mM): 68.5 NaCl, 50 KCl, 10 HEPES, 20 Glucose, 1 MgCl_2_, 2.5 CaCl_2_ at pH 7.4. The neurons were incubated with extracellular solution with or without high KCl for 30 min at 37°C. Some neurons were treated with 100 µM BAPTA-AM (Sigma) for 30 min before and during stimulation with high KCl extracellular solution. The extracellular solution was collected for ELISA analysis, centrifuged at 24,000 x g for 10 min and the supernatant was transferred to a new, low-protein-binding tube and stored on ice. The neurons were lysed with cold RIPA buffer described above. Plates with 96-wells (Corning 3925 “high-binding”) were prepared for ELISAs by coating each well with 50ul of 1.5ug/ml HT7 monoclonal antibody (Thermo Fisher) the night before use. Plates were sealed and incubated at 4°C, rocking, for ∼12 hr. The HT7 antibody was removed, wells were washed four times briefly (no incubation) with 300ul of DPBS, and then blocked in 300ul of 3% BSA/DPBS per well and incubated at room temperature, rocking, while samples were prepared as described above. A concentration series of seven Tau-441 protein standards were prepared for each plate. Each sample or standard (50uL) was added to individual blocked wells and incubated for 1 hr at room temperature with slow shaking. All samples and standards were run in triplicate. Then 50ul of Tau5-AP monoclonal antibody (1:500) in 0.3% BSA, 0.1% Tween-20, DPBS was added to each well and incubated overnight (∼12 hr) at 4°C with slow shaking. The next day, the plates were brought to room temperature (∼1hr). Wells were washed four times for 10 min in 300ul DPBS + 0.05% Tween-20 and tapped dry. Substrate CDP star was added (50ul per well) and the plates were incubated 30 min in the dark, followed by scanning on the plate reader. The CytoTox-ONE Membrane Integrity Assay (Promega) was performed according to the manufacturer’s instructions. The amount of secreted Tau was quantified from the percent of extracellular Tau detected out of the total amount of intracellular and extracellular Tau for each well of culture.

### Immunocytochemistry and Imaging

Human iPSC-derived neurons on coverslips were fixed in 4% paraformaldehyde in PBS for 15 min then washed three times for 5 min with PBS. The coverslips were then placed in blocking solution containing PBS, 0.1% Triton X-100, and 2% normal goat serum for 1 hour at room temperature. Primary antibodies diluted in blocking solution were added to the coverslips for 1 hour at room temperature then washed three times for 5 min with PBS. Primary antibodies used included mouse anti-flag (Sigma), anti-Munc18 (BD Biosciences), anti-Synaptotagmin 1 (Synaptic Systems) and rabbit anti-Synapsin (Cell Signaling Technology). The secondary antibodies diluted in blocking solution, including Alexa Fluor 488-conjugated anti-mouse and Alexa Fluor 546 anti-Rabbit (Life Technologies), were added to the coverslips for 1 hour at room temperature. Cy3-streptavidin (Jackson ImmunoResearch) was used for fluorescent labeling of biotinylated proteins. The coverslips were washed three more times for 5 min with PBS before mounting onto a slide with ProLong Gold Mountant (Thermo Fisher). Images were acquired with a laser scanning confocal microscope (LSM 700, Zeiss) using a 63x oil objective. The settings used for image acquisition were selected to keep the majority of brightest pixel intensities from reaching saturation. Maximum intensity projections were generated from confocal z-sections. Fluorescence intensity measurements were quantified using ImageJ software.

### Validation of TauWT-preferential interactors in postmortem human Alzheimer’s disease brain

Recent work identified six co-expressed protein modules (C1, C2, C3, C4, C8, and C10) that were significantly associated with Alzheimer’s disease progression using a multi-omics approach on multiple cohorts from the Accelerating Medicines Partnership – Alzheimer’s Disease (AMP-AD) consortium (Swarup et al., 2020). 29 out of the 108 proteins and 48 out of the 184 proteins, which preferentially interact with TauWT compared to TauP301L and TauV337M, respectively, were also in the six modules associated with AD. We determined if the number of intersecting proteins was significantly higher than expected using a permutation test (Efron and Gong, 1983). We considered all unique proteins identified by AP-MS from TauWT, TauP301L and TauV337M as the background set of proteins (N=1740). We randomly sampled 108 and 184 proteins, for TauP301L and TauV337M, respectively, from the background set of proteins and determined how many proteins overlapped with the six AD disease associated modules to create the null distribution. The p-value was calculated from the percentile of the null distribution that was ≥ 29 and ≥ 48 for TauP301L and TauV337M, respectively.

Label-free quantitative proteomic data from postmortem human brain tissues were downloaded for the Baltimore Longitudinal Study of Aging (BLSA) at Johns Hopkins University and Banner Sun Health Research Institute (Banner) from the Accelerating Medicines Partnership – Alzheimer’s Disease (AMP-AD) consortium (https://adknowledgeportal.synapse.org/) (Hodes and Buckholtz, 2016; Logsdon et al., 2019). The BLSA samples consisted of 97 samples from the dorsolateral prefrontal cortex (BA9 area) representing 15 controls, 15 AsymAD and 20 AD cases (Seyfried et al., 2017). The Banner samples were from prefrontal cortex of 30 controls, 28 mild cognitive impairment, 33 AsymAD and 98 confirmed AD cases (Swarup et al., 2020).

The label free quantitation intensities were log2 transformed and assessed for effects from biological covariates (diagnosis, age, gender) and technical variables (batch, brain bank). We used a linear regression model accounting for biological and technical covariates. The final model used was implemented in R version 3.6.1 (R Core Team, 2019) as follows:

lm(expression ∼ diagnosis + age + gender + batch + brain.bank.batch)

We evaluated the proteins that were TauWT preferential interactors compared to TauP301L (TauWT>TauP301L) and TauV337M (TauWT>TauV337M). We also evaluated the TauWT>TauP301L proteins and TauWT>TauV337M proteins that intersected with the mitochondrial C2 proteomic module (TauWT>TauP301L:C2 and TauWT>TauV337M:C2) from previous work (Swarup et al., 2020). In this work, the C2 proteomic module was significantly downregulated in Alzheimer’s disease compared to controls. For these protein sets, we considered their eigenprotein as the first principal component of their protein expression (Langfelder and Horvath, 2007; Zhang and Horvath, 2005).

We determined if the eigenprotein for TauWT>TauP301L, TauWT>TauV337M, TauWT>TauP301L:C2 and TauWT>TauV337M:C2 was significantly different from disease diagnosis (AD, mild cognitive impairment, AsymAD) versus control using the Wilcoxon rank sum test (Wilcoxon, 1945). Eigenproteins were correlated with neuropathological scores (CERAD, Braak) using Pearson correlation (Pearson, 1931).

### Mitochondrial respiration

Oxygen Consumption Rate (OCR) from live human iPSC-derived neurons was measured using a Seahorse XFe-96 Analyzer (Agilent). 30,000 human pre-differentiated neurons were seeded per well in Poly-D-Lysine + Laminin-coated 96-well Seahorse plate and differentiated for the indicated time (2 or 4 weeks) before performing the OCR measurements. 1 hour before the Seahorse experiment, neurons were washed and incubated with the assay media (Agilent Seahorse XF Base Medium (Agilent, Cat# 103335-100) supplemented with 20mM Glucose, 1.2mM Glutamine, 1mM Sodium Pyruvate, pH=7.4) in a non-CO_2_ incubator. Oligomycin (final concentration=1uM, Fisher Scientific, Cat# 49-545-510MG), FCCP (final concentration=1uM, Fisher Scientific, Cat# 04-531-0) and a mixture of Rotenone (final concentration=0.5uM, Fisher Scientific, Cat# 36-165-0) and Antimycin A (final concentration=0.5uM, MilliporeSigma, Cat# A8674) were sequentially injected into the media to modulate distinct components of the ETC and reveal key parameters of mitochondrial bioenergetics. After the OCR measurements were obtained, neurons were incubated with Calcein-AM (Invitrogen, Cat# C3099) for 30 min, and imaged using ImageXpress® Pico Automated Cell Imaging System (Molecular Devices). OCR was normalized to live neurons using Calcein-AM staining intensity.

### Mitochondrial membrane potential measurement using TMRM

iPSC-derived neurons were cultured in PDL-coated 96-well plate (Cat#354640, Corning) for 4 weeks. The day of the experiment, neurons were washed twice in assay medium containing 50% phenol-free Neurobasal-A media plus 50% phenol-free DMEM/F12 supplemented with N2 supplement (0.5X), B27 supplement (0.5X), Glutamax (0.5X), Non-Essential Aminoacids (1X), BDNF (10 ng/mL) and NT3 (10 ng/mL). Neurons were incubated with assay media containing 10nM Tetramethylrhodamine, methyl ester (TMRM) (Cat# I34361, Invitrogen) and 1ug/ml Hoechst 33342 (Cat# H3570, Invitrogen) at 37°C for 30 minutes. Neurons were imaged at 37°C and 5% CO2 using the KEYENCE BZ-X700 microscope.

## SUPPLEMENTARY FIGURES

**Figure S1.**
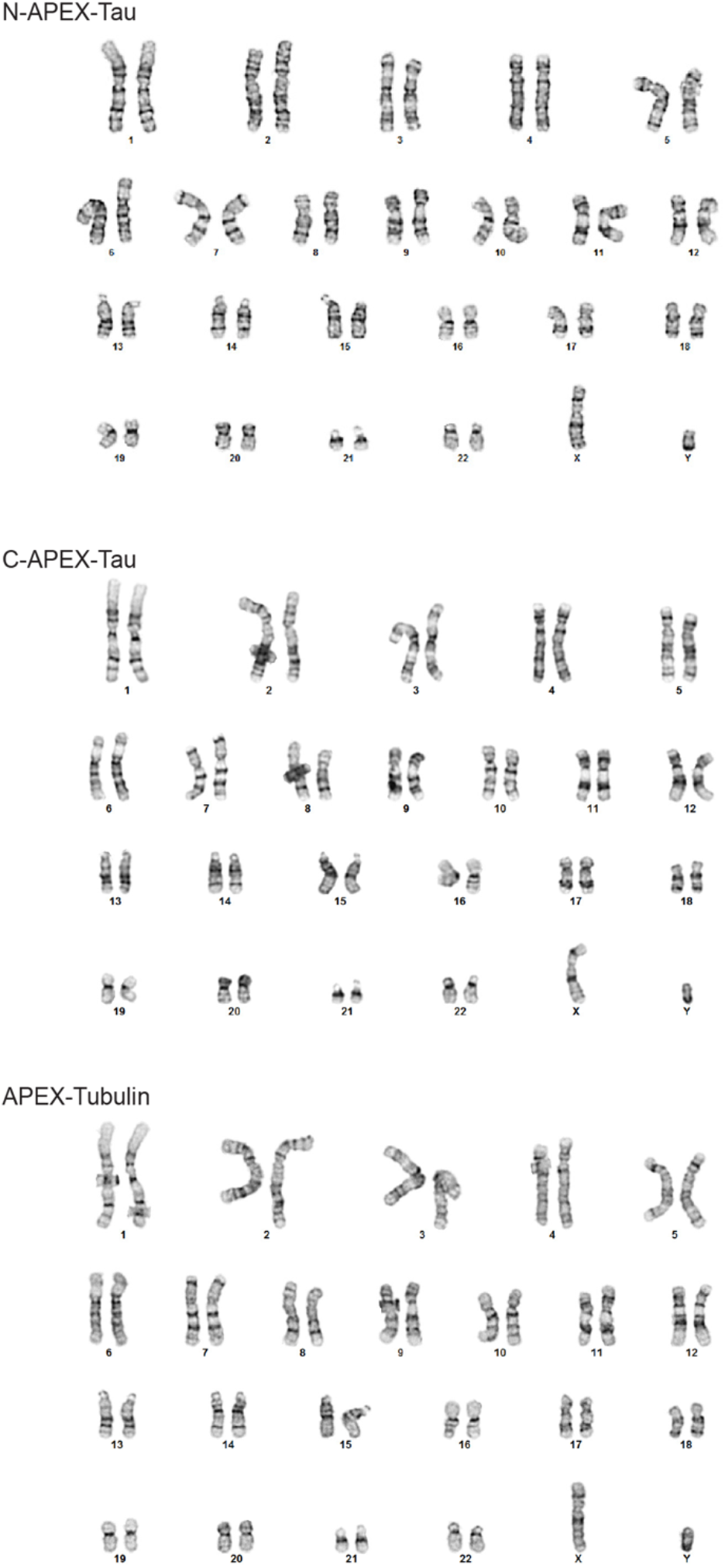
Karyotyping analyses of human iPSCs related to Figure 1. The isogenic iPSCs clones generated for these studies, including N-APEX-tau, C-APEX-tau and APEX-α tubulin iPSCs had normal Karyotypes.

**Figure S2.**
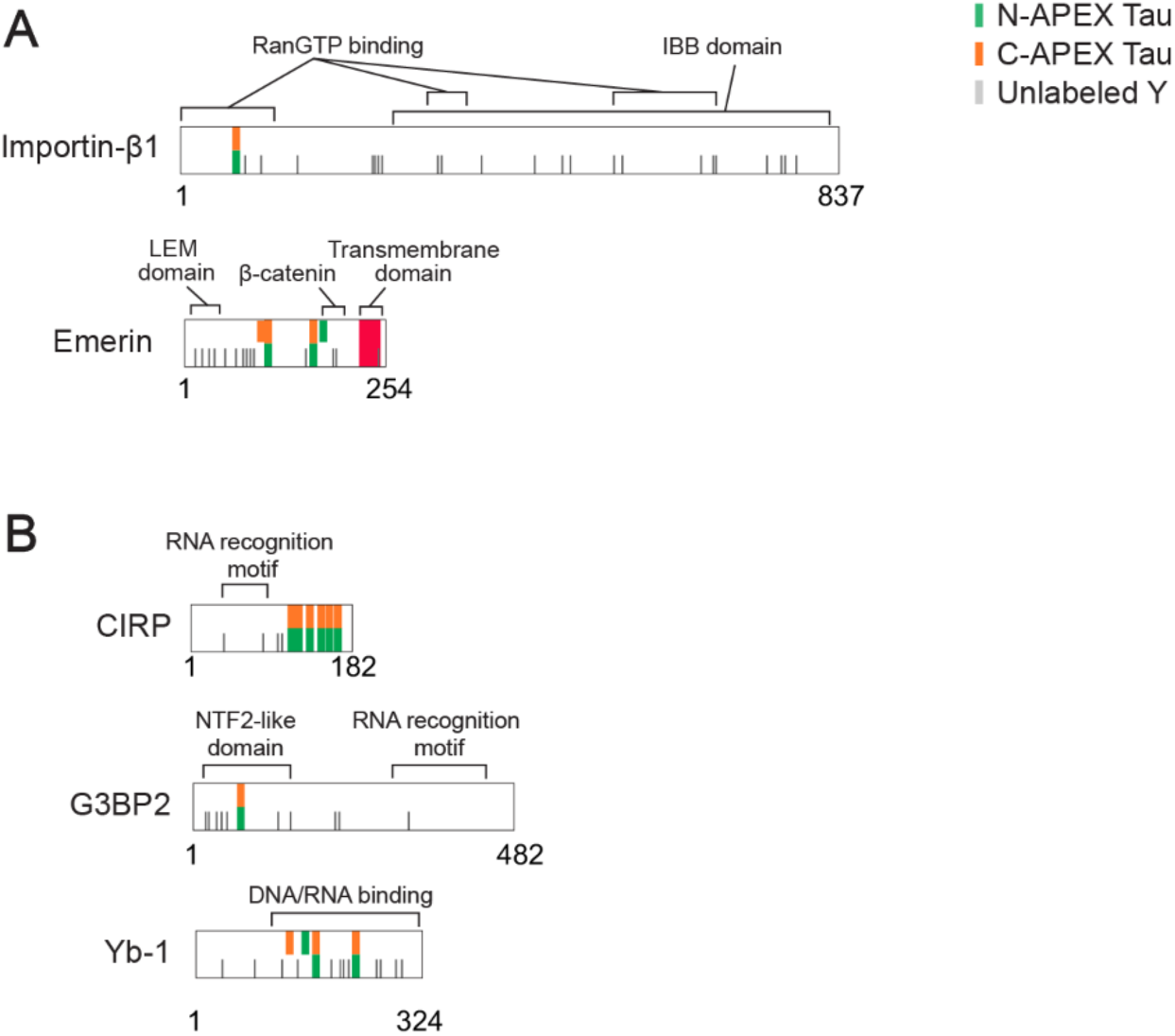
Biotinylation sites detected on proteins localized at the nucleus and bound to RNA or DNA. **(A)** Biotinylated tyrosine residues detected with N-APEX Tau (green) and C-APEX Tau and unlabeled tyrosines (grey) mapped onto nuclear proteins. **(B)** Biotinylated tyrosine residues and unlabeled tyrosines mapped onto RNA- and DNA-bindingproteins.

**Figure S3.**
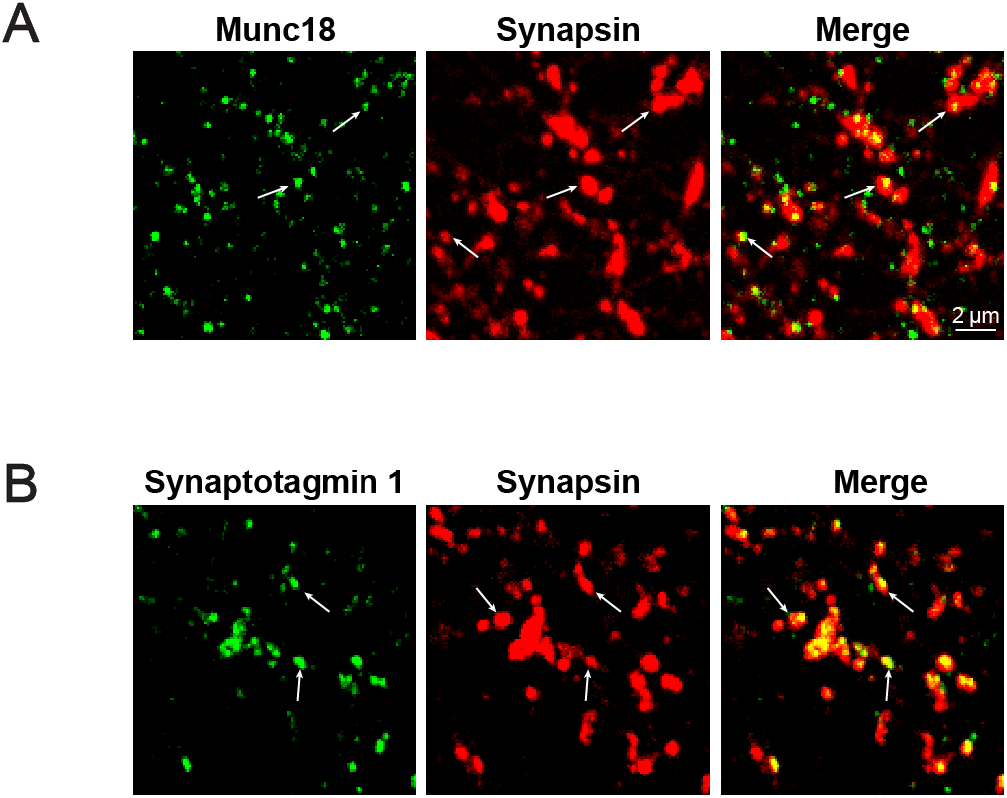
Munc18 and Synaptotagmin 1 are localized at presynaptic terminals in human neurons related to Figures 3 and 4. Representative confocal images of immunostaining in mature wildtype human iPSC-derived neurons showing **(A)** co-localization of Munc18 (green) and Synapsin (red), a marker of presynaptic terminals as well as **(B)** co-localization of Synaptotagmin 1 (green) with Synapsin (red).

**Figure S4.**
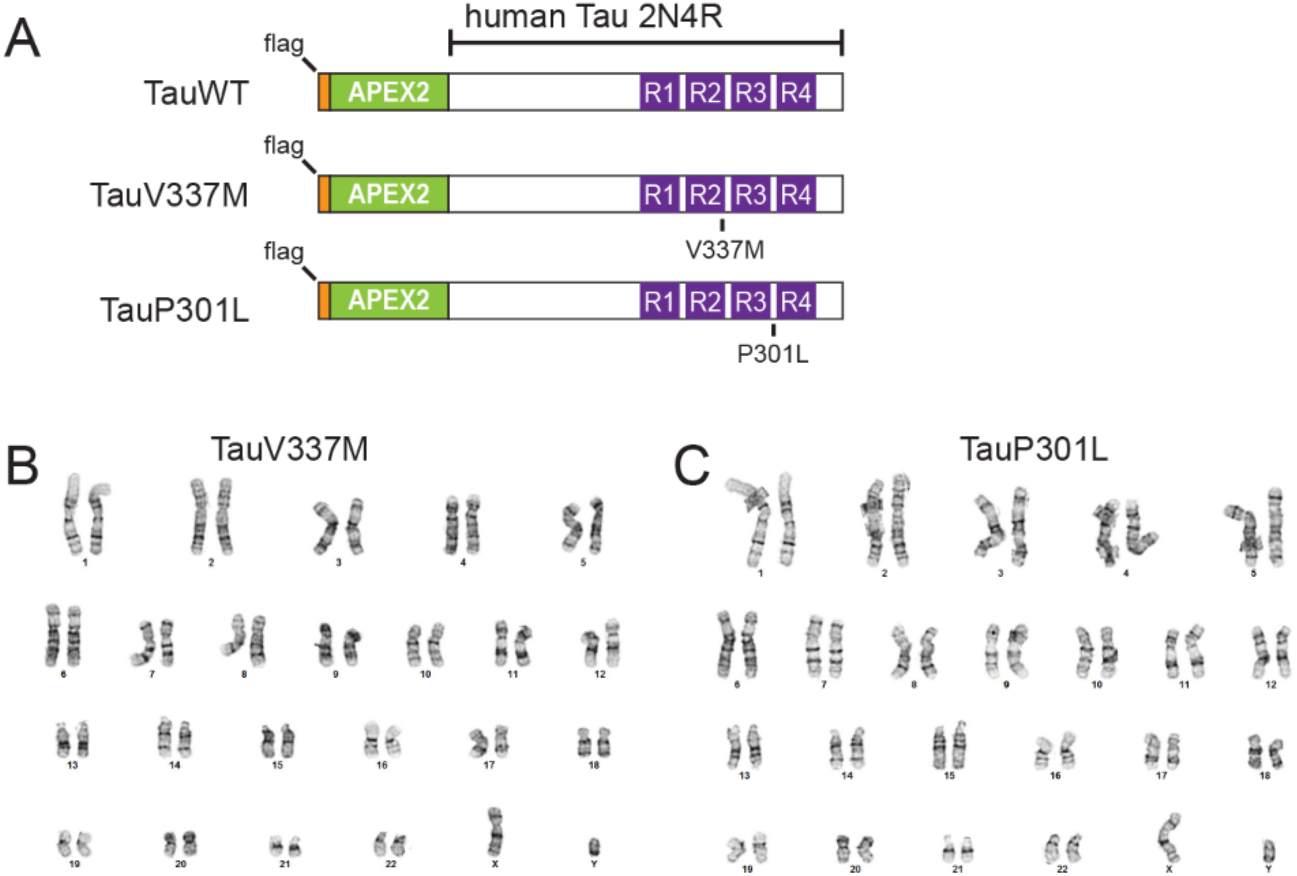
Karyotyping analyses of FTD mutant Tau human iPSCs related to Figure 5. **(A)** Design of the wildtype (WT), V337M and P301L APEX-flag tagged full-length human Tau (2N4R) constructs expressed in human iPSC-derived neurons. (Band C) Karyotype analyses of isogenic iPSC clones with TauV337M and TauP301L were normal. The isogenic TauWT clone karyotyping results are shown in Figure S1.

**Figure S5.**
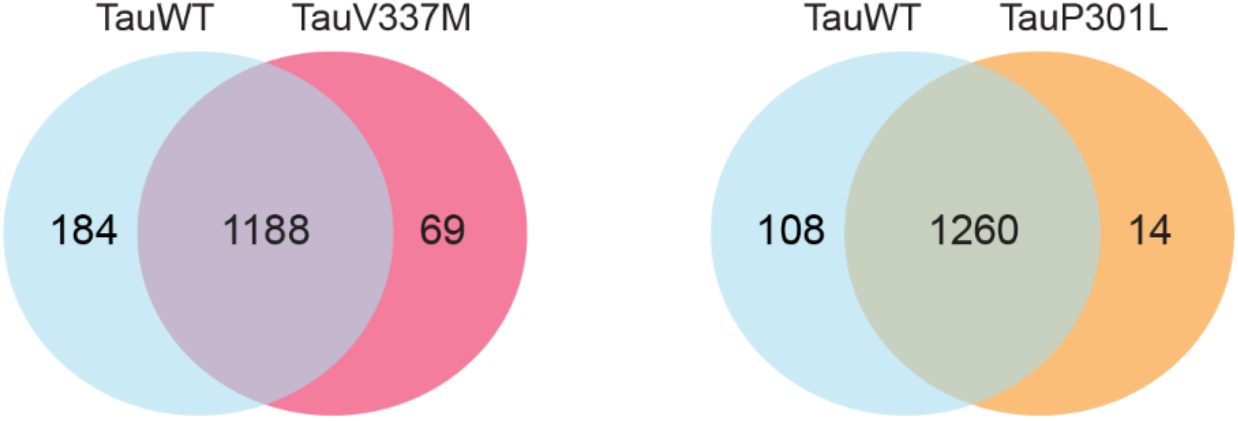
Comparison of protein-protein interactions identified for TauWT and Tau with FTD-causing mutations related to Figure S and Table SS. Venn diagrams of AP-MS results from human neurons indicating the overlap of proteins bound to Tau WT and TauV337M (left) or the overlap between TauWT and TauP301 L (right). See also Tables S5.

**Figure S6.**
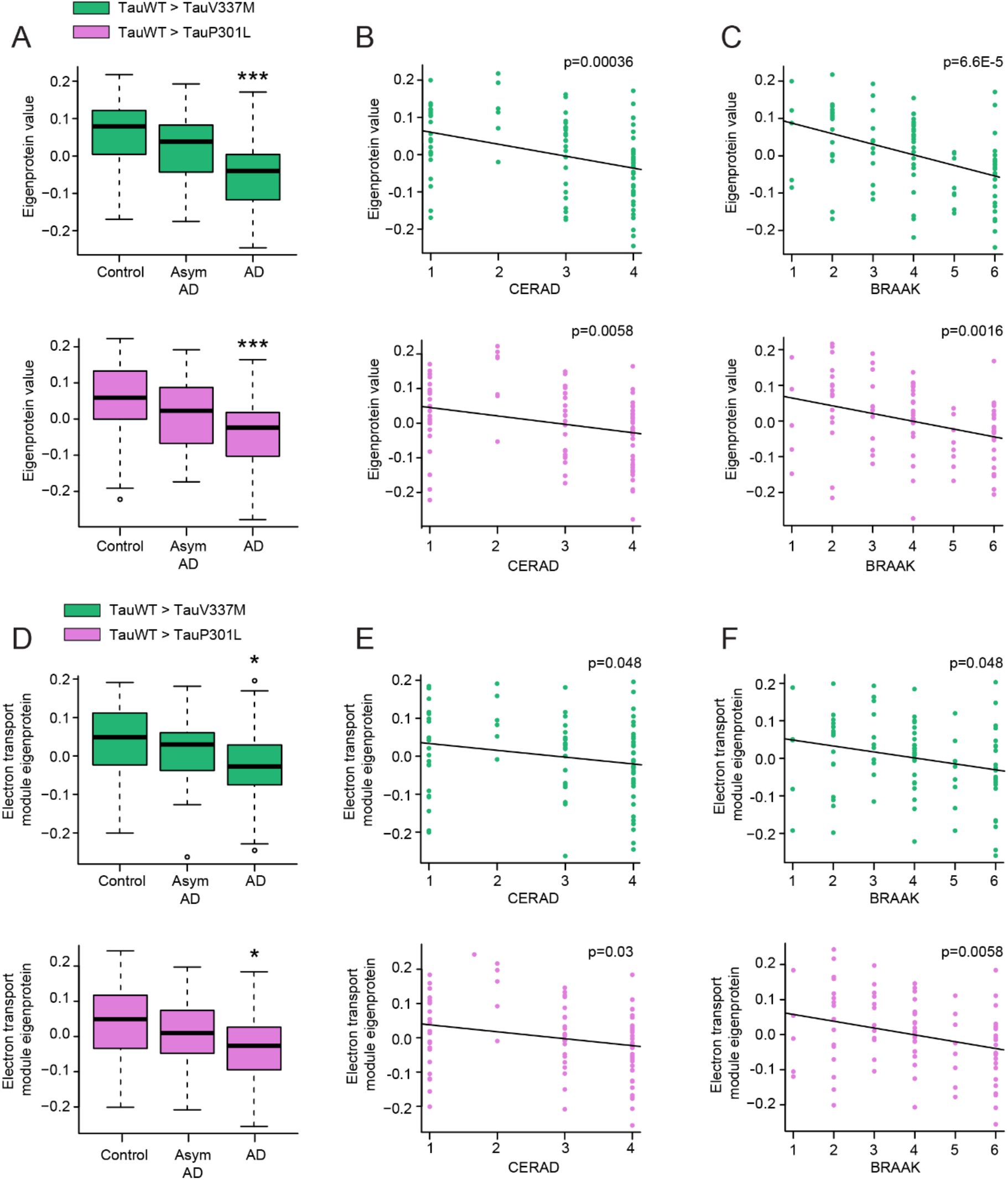
BLSA cohort module analyses of the levels of TauWT-preferential interactors in human brain related to Figure 8. **(A-C)** Plots showing TauWT-preferential interactor eigenprotein trajectory with AD diagnosis **(A),** CERAO **(8),** and BRAAK **(C)** scores in BLSA cohort. TauWT-preferential interactors as compared to TauV337M and P301Lare colored as green and pink, respectively. ***p < 0.005. **(D-F)** Plots showing C2-mitochondrial module TauWT-preferential interactor eigenprotein trajectory with AD diagnosis **(D),** CERAO **(E),** and BRAAK **(F)** scores in BLSA cohort. TauWT-preferential interactors as compared to TauV337M and P301L are colored as green and pink, respectively. *p < 0.05.

